# SUBTLE DIFFERENCES IN FINE POLYSACCHARIDE STRUCTURES GOVERN SELECTION AND SUCCESSION OF HUMAN GUT MICROBIOTA

**DOI:** 10.1101/2022.10.04.510853

**Authors:** Tianming Yao, Dane G. Deemer, Ming-Hsu Chen, Bradly L. Reuhs, Bruce R. Hamaker, Stephen R. Lindemann

## Abstract

Dietary fibers are fermented in the human gut and are known to modulate microbiome composition and metabolic function, but few studies have explored to what extent the small variations in complex fiber structures impact community assembly, microbial division of labor, and organismal metabolic responses across individuals’ microbiome structures. To test the hypothesis that subtle linkage variations in chemical structures of polysaccharides afford different ecological niches for distinct communities and metabolism, we employed a 7-day *in vitro* sequential batch fermentation with fecal inocula from individual donors and measured microbial responses using an integrated multi-omics approach. We fermented two sorghum arabinoxylans (SAXs) as model complex polysaccharides, with fecal microbiota from three donors and an artificially high diversity mix of all three. Although differences in sugar linkage profiles across SAXs were subtle, surprisingly, consortia fermenting different AXs revealed distinct species-level genomic diversity and metabolic outcomes with nearly-identical strains on each polysaccharide across inocula. Carbohydrate active enzyme (CAZyme) genes in metagenomes revealed broad AX-related hydrolytic potentials; however, CAZyme genes enriched in different AX-fermenting consortia were specific to SAX type and displayed various catabolic domain fusions with diverse accessory motifs, suggesting they may be functionally degenerate and this degeneracy may relate to fine substrate structure. These results suggest that fine polysaccharide structure exerts deterministic selection effect for distinct fermenting consortia, which are present amongst unrelated individuals.

## INTRODUCTION

The human gut microbiota play an integral role in health and disease [1, 2]. Colonic bacteria vigorously synthesize bioactive metabolites, many of which prevent colon diseases (e.g., inflammation, colitis, and cancers) and beneficially modulate host metabolism [3–6]. Manipulation of gut microbiota function can be achieved by consumption of dietary fibers, which are prime substrates for saccharolytic microbial fermentation. Previously, we demonstrated that microbiota competing for carbohydrate polymers are strongly governed by substrate chemical complexity [7]. However, information is still limited on how microbiota respond metabolically to discrete fibers across idiosyncratic gut microbiomes and how fine structural variations of complex polysaccharides impact gut community assembly and division of labor (DOL) among members.

Recently, mounting evidence has suggested tight relationships among some complex glycans and fermenting microbes. Fibers with more complicated chemical structures were more specific and predictable in targeting particular gut microbiota [8, 9], since diverse chemical linkages and glycosyl units of complex glycans likely offer multiple ecological niches for specialist microbes. The decomposition of complex polymers further structures the downstream assembly of the community with diverse oligomeric structures, which select for secondary consumers and influence the metabolic exchanges by the members [10]. In contrary, simple fibers are widely accessible to gut organisms because few enzymes are required to degrade them. Thus, they may therefore not support niche differentiation, instead promoting strong competition which may lead over time to competitive exclusion and concomitant reductions in diversity, stability, and possibly metabolic function [7].

Complex polysaccharides may maintain diversity via DOL in multiple ways. Because complex polysaccharides molecules are too large to be directly transported into microbial cells, extracellular hydrolytic enzymes are needed to initiate degradation. These carbohydrate active enzymes (CAZymes) are highly specific to hydrolysis of cognate polysaccharide ligands, typically at the level of disaccharide bonds [11]. Consequently, a complex fiber with diverse glycosidic linkages requires multiple distinct CAZymes for complete degradation, meaning that diverse microbial members encoding different hydrolytic genes may be required. Microorganisms with complementary suites of CAZymes or compatible regulation of CAZyme genes may divide labor in hydrolyzing substrates, performing cooperative degradation [12]. However, limited mechanistic understanding exists regarding how fiber chemical structures affect the niche structure of fermenting communities. Here, we hypothesized that 1) complex glycan structures exert strong deterministic selection in community assembly and will recruit similar fermenting consortia across distinct individuals’ microbiota, and 2) subtle linkage variations in the fine chemical structure of a complex fiber afford different ecological niches that will select for distinct communities and metabolism.

To test these hypotheses, we performed sequential *in vitro* batch fermentation of two arabinoxylans (AXs) with subtly differing linkage profiles with human fecal microbiota under high dilution pressure, selecting for consortia of organisms able to grow most rapidly on each model fiber. AXs are abundant hemicelluloses in cereals and have complex fine chemical structures (>12 different carbohydrate linkages) and large molecular weights (>10^6^ Daltons) that vary across plant sources. We report that consortia selected across donor inocula converged rapidly in complex carbohydrate fermentations, forming reproducible, stable, and diverse communities whose compositions were driven by fine AX chemical structures. Further, the CAZyme genes enriched in each stable community were driven by substrate fine structure, suggesting fitness differences across strain-level genotypic variants that are specific to fibers. Together, our data suggest that 1) complex fibers exert strong selective pressure on fermenting communities, even across divergent inocula, and 2) even subtle variances of fine polysaccharide structures select for consortia varying in diversity and composition and with distinct metabolic function.

## RESULTS

### Subtly different AX chemical structures select for distinct consortia with diversity related to fiber complexity

The chemical structures of two sorghum-derived AXs varied subtly in linkage composition and fine structures (Fig. 1A, B and Table 1). The arabinose:xylose ratio was significantly higher (*p* < 0.05) in red sorghum bran AX (RSAX) than white sorghum bran AX (WSAX), suggesting more branching in RSAX. From linkage profiles (Table 1), different AX backbone substitution patterns were observed. The unsubstituted backbone linkage “→4(Xyl*p*)1 →” was not significantly different among AXs, whereas the proportion of singly-substituted backbone “→4(Xylp)1 →3↑” and the di-substituted backbone “→4(Xylp)1 →3↑↑2” was greater for WSAX and RSAX, respectively. Regarding side chain linkages, “→3(Ara*f*)1 →” was significantly elevated in WSAX whereas “→5(Ara*f*)1 →3↑↑2” was higher in RSAX. Although both linkages may compose complex branches, the linkage “→5(Ara*f*)1 →3↑↑2” generally forms a more complex side chain with at least three sugar units [13], suggesting overall more complex branching in RSAX.

**Table 1.** Glycosidic linkage profiles (mol %) of red and white SAX ^A, B^

|  | Linkage indicated | RSAX (mol %) | WSAX (mol %) |
| --- | --- | --- | --- |
| Xylan backbone linkages | →4(Xylp)1→ | 8.00 ± 1.25 <sup>a</sup> | 9.37 ± 0.74 <sup>a</sup> |
|  | →4(Xylp)1→3↑ | 15.21 ± 1.57 <sup>b</sup> | 17.77 ± 0.30 <sup>a</sup> |
|  | →4(Xylp)1→3↑↑2 | 17.53 ± 1.78 <sup>a</sup> | 14.62 ± 0.35 <sup>b</sup> |
| Side-chain linkages | (Araf)1→ | 20.91 ± 0.87 <sup>a</sup> | 19.30 ± 0.98 <sup>a</sup> |
|  | (Xylp)1→ | 1.01 ± 0.13 <sup>a</sup> | 1.04 ± 0.13 <sup>a</sup> |
|  | (Manp)1→ | 2.48 ± 0.30 <sup>a</sup> | 2.69 ± 0.10 <sup>a</sup> |
|  | (Galp)1→ | 6.65 ± 0.00 <sup>b</sup> | 7.59 ± 0.00 <sup>a</sup> |
|  | (Glc p)1→ | 1.91 ± 0.31 <sup>a</sup> | 1.77 ± 0.52 <sup>a</sup> |
|  | →2(Araf)1→ | 1.5 ± 0.29 <sup>a</sup> | 2.01 ± 0.17 <sup>a</sup> |
|  | →3(Araf)1→ | 1.98 ± 0.02 <sup>b</sup> | 2.43 ± 0.01 <sup>a</sup> |
|  | →5(Araf)1→ | 1.81 ± 0.03 <sup>a</sup> | 2.60 ± 0.66 <sup>a</sup> |
|  | →5(Araf)1→3↑ | 2.00 ± 0.66 <sup>a</sup> | 1.04 ± 0.11 <sup>a</sup> |
|  | →5(Araf)1→3↑↑2 | 10.99 ± 0.14 <sup>a</sup> | 7.10 ± 0.72 <sup>b</sup> |
|  | →4 (Manp)1→ | 1.97 ± 0.30 <sup>a</sup> | 2.06 ± 0.10 <sup>a</sup> |
|  | →4 (Glc p)1→ | 6.04 ± 0.31 <sup>a</sup> | 8.61 ± 0.52 <sup>a</sup> |
| Degree of branches |  | 1.23 ± 0.10 <sup>a</sup> | 1.13 ± 0.01 <sup>a</sup> |
| Branch complexity |  | 38.35 ± 1.00 <sup>a</sup> | 38.93 ± 1.78 <sup>a</sup> |

<sup>A</sup> Values are expressed as a proportion of all partially methylated alditol acetates present.<sup>B</sup> Different letters in each row indicates significant differences (p < 0.05) between samples

**Figure 1:**
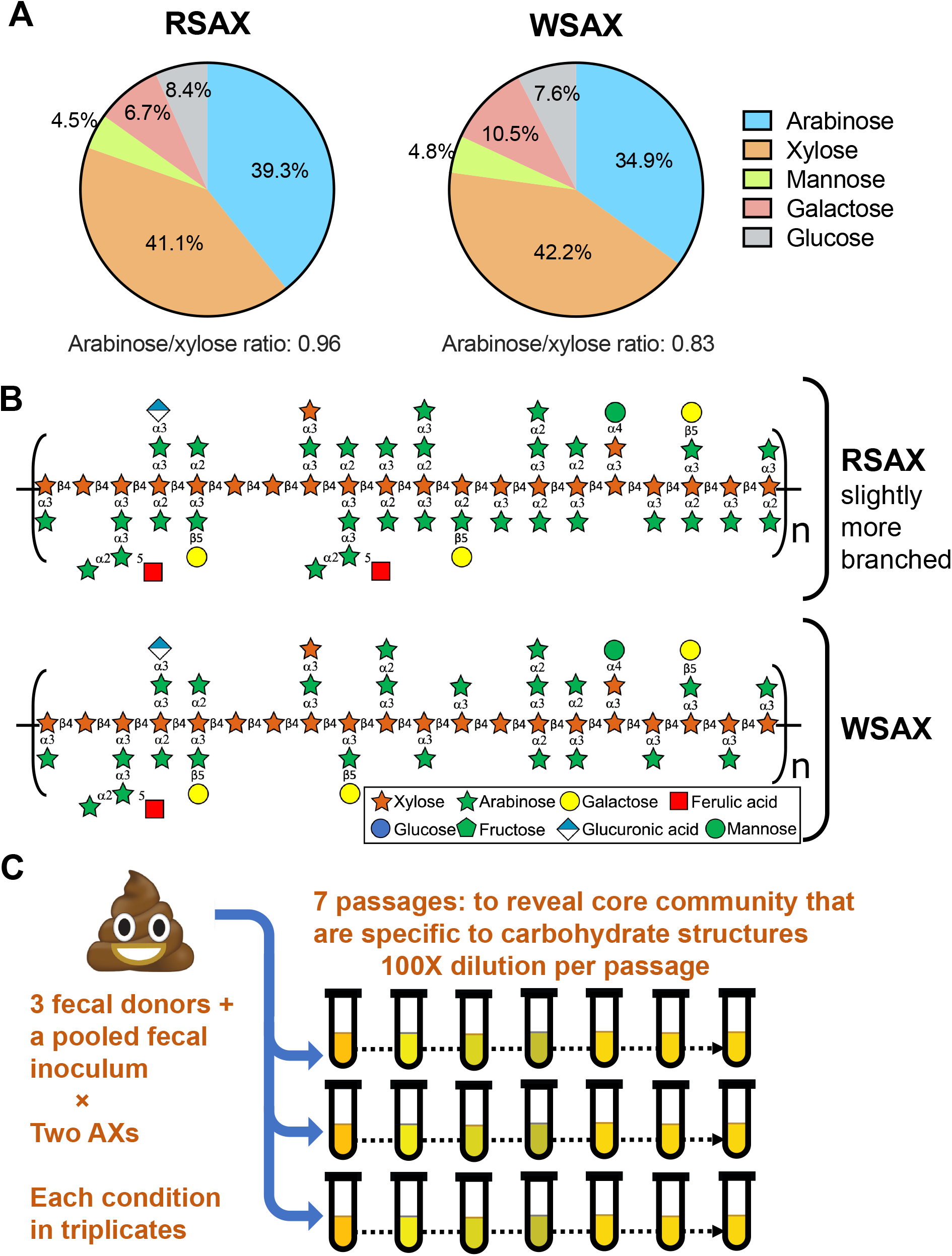
Monosaccharide composition (A) of two purified sorghum arabinoxylan substrates used in sequential fermentations (mol %). Schematic linkage model (B) of RSAX and WSAX polysaccharide structures. Experimental design of a 7-day sequential passage fermentation (C), revealing consortia evolution on substrate affinity. The 1:100 dilution between two passages forms bottleneck effects, which screen out irrelevant (slow-growing) bacteria and retain the substrate-specific core community.

Succession trajectories among triplicated lineages were similar over the 7-day sequential passage fermentation (one representative lineage is presented in Fig. 2 and all lineages in Fig. S1), suggesting that AXs exerted strong deterministic forces on community assembly. Although the inoculum governed community structure in early passages, cultures on distinct SAXs converged rapidly and strongly. Dominance of the same operational taxonomic units (OTUs) was found across donors, which were classified within genera *Bifidobacterium* (Otu003), *Bacteroides* (Otu002), *Phocaeicola* (Otu004), and *Agathobacter* (Otu001). Some dominant OTUs at passage 7 were undetectable in initial fecal samples at our normalized sequencing depth (e.g., Otu001 was non-detectable in the lowest initial diversity inoculum (Donor 2) but ended with 25.9% average read abundance in the final WSAX passage) (Fig. 2B). Interestingly, these initially rare taxa displayed some evidence of stochasticity in consortial assembly. For example, two replicates of Donor 2 (which had initially small populations of phylum Bacillota OTUs) displayed sizable final populations of Otu001 in WSAX consortia, however, in the third lineage, Otu001 reads were not observed in any intermediate or final culture (Fig. S1A). Conversely, only one of three lineages of Donor 2 RSAX exhibited detectable populations of Otu001. Artificially high initial diversity (pooled inocula) appeared to enhance the determinism of fiber structure on community succession, displaying smaller inter-lineage differences in community structure.

**Figure 2:**
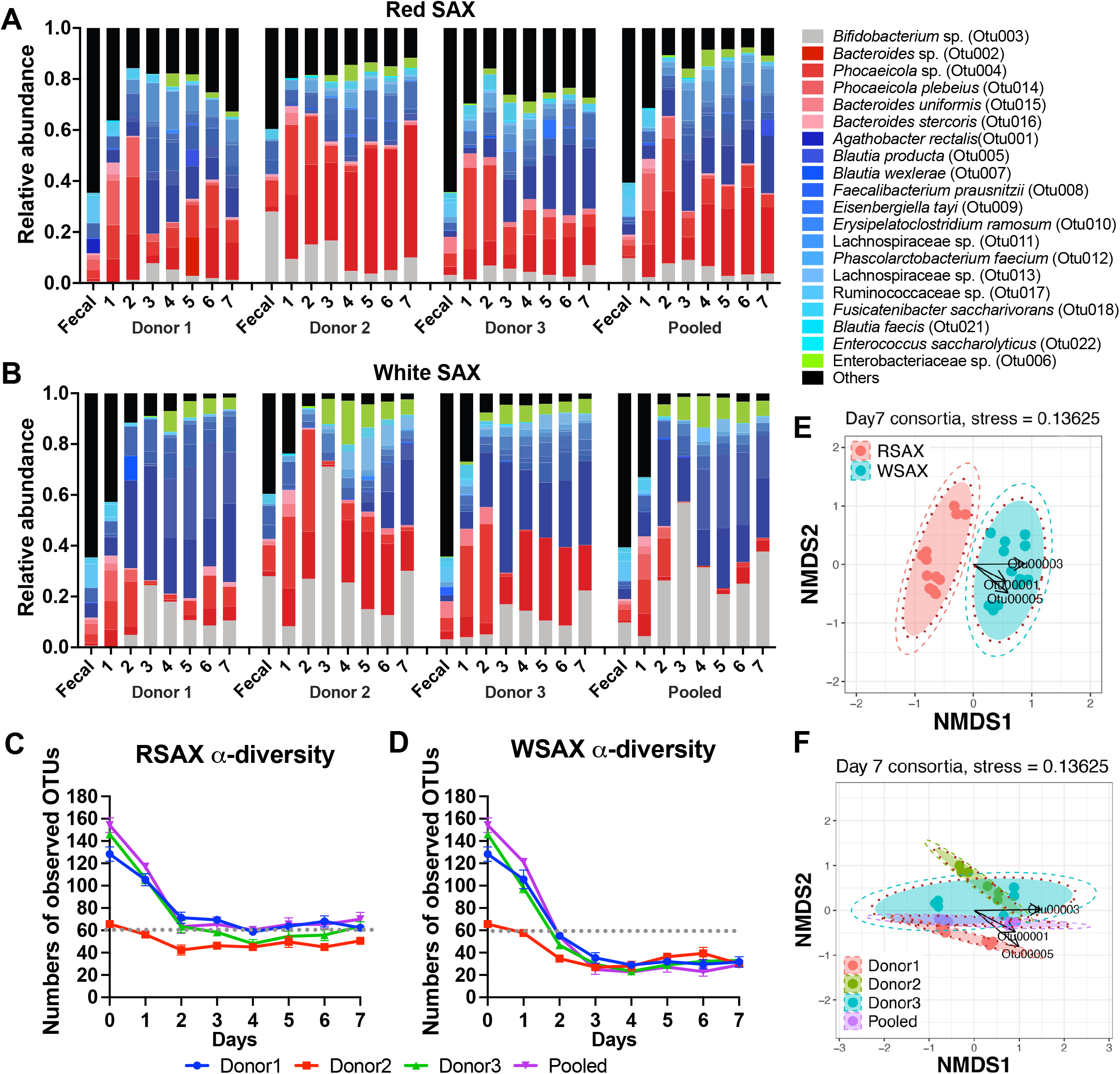
Microbial community structure over 7 passages in RSAX and WSAX fermentations revealed by 16S rRNA gene amplicon sequencing. Changes of community structures from one lineage of each inoculum over 7 days are shown in sequential cultures on RSAX (A) and WSAX (B). Each shade represents a distinct OTU, which are colored by phylum: Pseudomonadota (green), Bacillota (blue), Bacteroidota (red), and Actinomycetota (gray). Rare OTUs (below 0.2% relative abundance) are grouped in Others (black). Alpha-diversity curves of evolving communities on RSAX (C) and WSAX (D). Non-metric multi-dimensional scaling (NMDS) plots on day-7 consortia, grouped by substrate differences (E) or inter-donor differences (F). The outer circles with dashed line indicate separation of group clusters at the 99% confidence interval.

Amplicon data suggested that the more-complex RSAX fiber sustained a higher microbial diversity than WSAX, particularly with respect to genus *Bacteroides* (Fig. 2C, D). Both AXs’ communities revealed initially steep losses in α-diversity curves over dilution which stabilized at later passages, suggesting slow-growing species were rapidly diluted out as core substrate-consuming consortia coalesced. Bacterial single copy gene (BSCG) counts of passage 7 shotgun metagenomic assemblies also supported higher α-diversity in RSAX consortia (Fig. S2). Ordination plots of final consortia community structure (Fig. 2E, F) and community structure variation over the 7 passages (Fig. S3A) revealed a significant separation of two AX-fermenting consortia, suggesting strong convergence around polysaccharide structure instead of donor effects (Fig. S3B-D).

### RSAX and WSAX were fermented to different metabolic outcomes

To determine whether differences in polysaccharide structure and fermenting consortium composition impacted metabolic function, we measured gas production (Fig. 3A), pH (Fig. 3B) and SCFA production (Fig. 3C & Fig. S4A) at every passage. Gas production was insignificantly different across substrates and inocula except that it was elevated in WSAX-Donor 3. Cultures increased in terminal pH until day 3 and stabilized by day 5, suggesting consortia had reached functional stability by that point. RSAX cultures generally ended at higher pH than WSAX cultures for each donor, although RSAX fermentation evolved significantly more total SCFAs than WSAX (except Donor 1, Fig. 3C), indicating differences in metabolite production.

**Figure 3:**
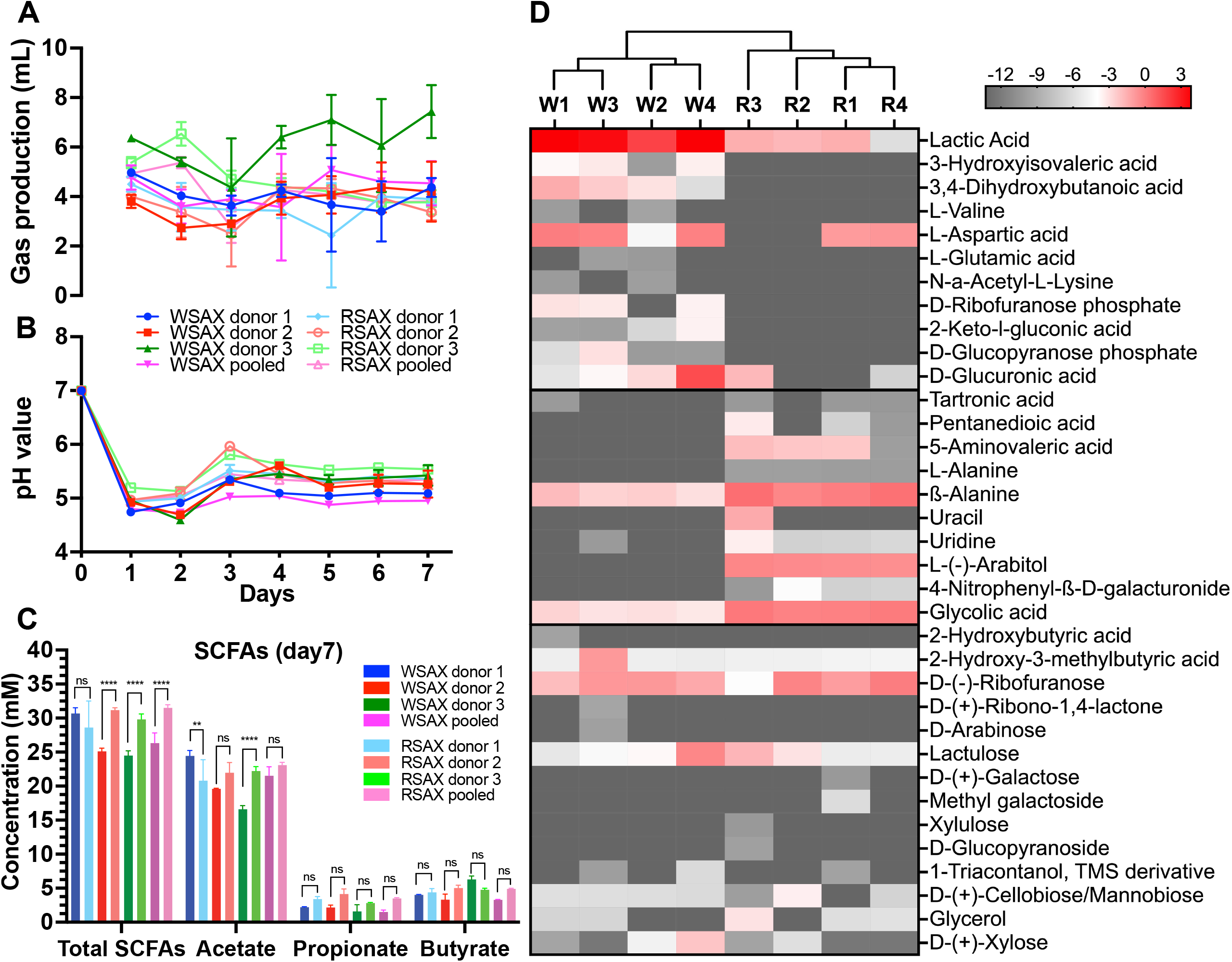
Total gas production (A) and pH curves (B) from sequential batch cultures. Total short chain fatty acids (SCFAs), acetate, propionate and butyrate production comparison across substrates at day 7 (C). Statistically significant differences between substrates were calculated by Tukey’s multiple comparisons test at *p* < 0.05. Symbol style: nonsignificant (ns), 0.0332 (*), 0.0021 (**), 0.0002(***), <0.0001(****). Heatmap of untargeted polar metabolomics (D). Average peak areas of triplicates were normalized by percent of total detected areas and with log2-transformation for display. Red and grey colors represent high and low abundance of metabolites, respectively.

To determine whether broader metabolic outcomes were impacted by SAX structure-consortium interactions, we performed untargeted polar metabolomics on final cultures (Fig. 3D), and targeted HPLC analysis on supernatant concentrations of lactate and propanediol (Fig. S4C). RSAX fermentations produced significantly higher (*p* < 0.05) beta-alanine and glycolic acid than WSAX ones, and some metabolites were only found in RSAX cultures (e.g. pentanedioic acid, 5-aminovaleric acid, L-alanine, and L-arabitol). Conversely, WSAX fermentations produced significantly higher (*p* < 0.05) lactic acid, L-aspartic acid, and D-glucuronic acid. Moreover, 3-hydroxyisovaleric acid, 3,4-dihydroxybutanoic acid, L-valine, L-glutamic acid, N-a-acetyl-L-lysine, D-ribofuranose phosphate, 2-keto-l-gluconic acid, and D-glucopyranose phosphate were exclusively detected in WSAX supernatants. HPLC analysis further quantified significantly higher (*p* < 0.01) propanediol associated with RSAX supernatants and large amounts of lactic acid (2-4 mM, Fig. S4C) in WSAX fermentations, likely driving lower pH in WSAX cultures despite their low SCFA production. Such explicit differences in metabolite composition among substrates revealed that a subtle AX structure difference strongly governed metabolic outcomes of fermentation, even given divergent donor inocula. However, abundances of several metabolites, such as glycerol and D-xylose, varied across different donors’ consortia fermenting identical SAX, suggesting some stochastic influence of consortium membership.

### Fiber structure selected nearly-identical strains across donor communities, but strain penetrance was species-dependent

Reconstruction of metagenome-assembled genomes (MAGs) from final consortia provided species- and strain-level resolution that is not differentiable using 16S amplicons. For example, only one OTU from genus *Bifidobacterium* was identified from 16S amplicons, although we recovered MAGs from four bifidobacterial species, namely *B. longum, B. dentium, B. pseudocatenulatum* and *B. adolescentis* (Table S1). Interestingly, these four bifidobacterial species co-existed in one consortium (WSAX-pooled), suggesting the hypothesis that they may coexist by dividing labor on WSAX-derived substrates. Since strains of the same species can vary significantly in their genome content and metabolic function, we performed strain-level examinations of gene content differences across strains from different donors and related those differences to prevalence in consortia fermenting each SAX (especially pooled inocula, where each donors’ strains were forced into competition with one another).

Surprisingly, in many cases, nearly-identical MAGs across donors’ consortia were observed (Fig. 4). For example, *A. rectalis* strains across donors’ consortia clustered tightly together except for strains in RSAX-pooled and WSAX-Donor-3. Within genus *Bifidobacterium, B. longum* was universally found in all 8 consortia and formed 3 clades of nearly-identical strains: *B. longum* MAGs from Donor 2 (RSAX and WSAX), Donor 3 (RSAX), and pooled (WSAX) clustered together, Donor 3 (WSAX) and pooled (RSAX) formed a second cluster, and Donor 1 strains (both SAX types) composed the third. *B. adolescentis* MAGs were identified with two clades of nearly identical strains in RSAX (Donor 2 & 3) and WSAX (Donor 2 & pooled). These data suggested that identical strains were present even across different individuals and successfully competed in SAX-consuming consortia, but not necessarily in substrate-specific ways. MAGs from genus *Bacteroides*, another key SAX-degrading genus, showed evidence of donor-specificity and increased inter-strain competition (Fig. S5), suggesting that different *Bacteroides* spp. may be competitive under differing initial inoculum communities.

**Figure 4:**
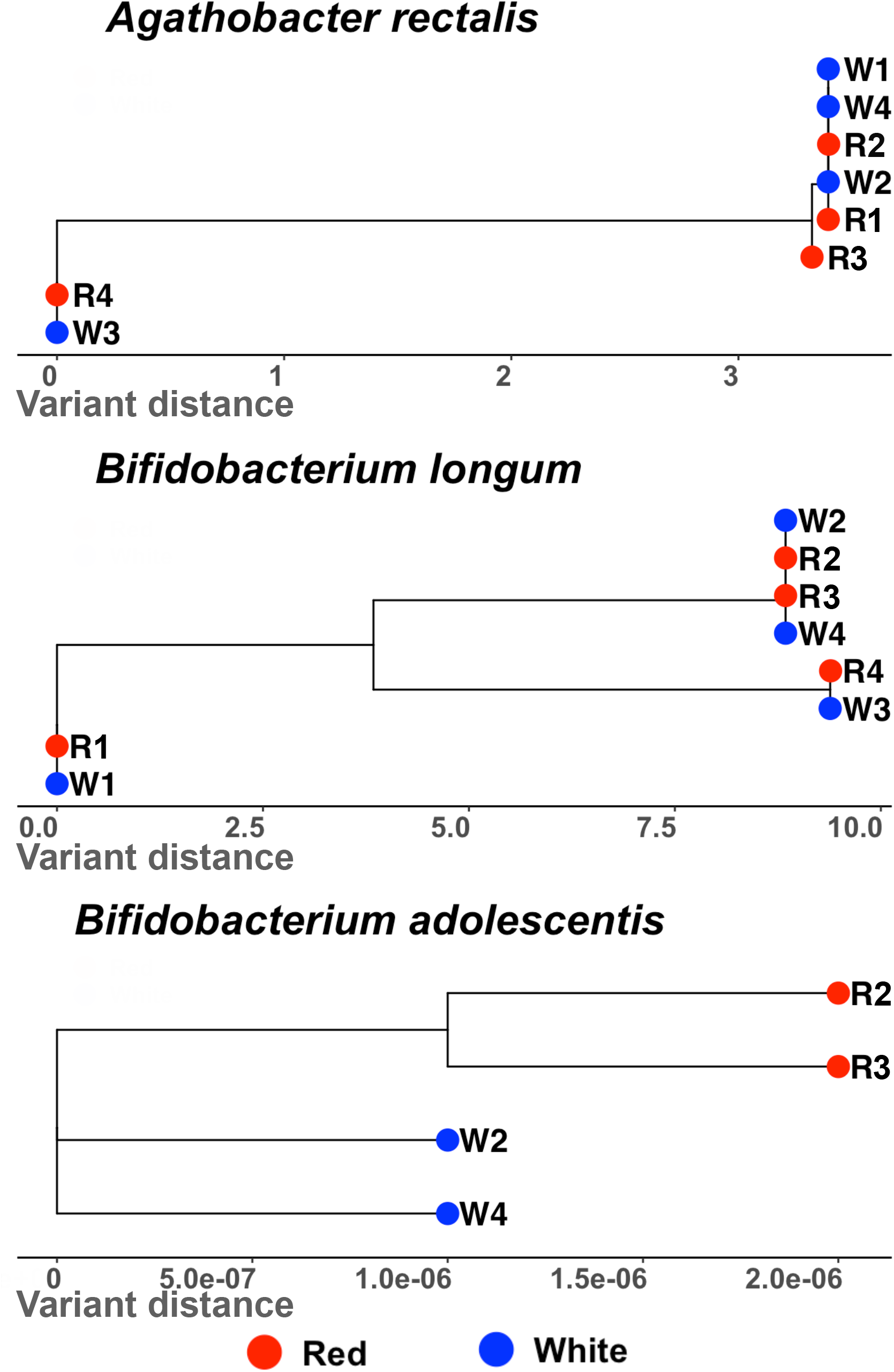
Strain level comparison of *Agathobacter rectalis* and bifidobacterial MAGs from day-7 metagenomes, identified by StrainPhlan. Label “W4” and “R4” denote WSAX-pooled and RSAX-pooled samples, respectively. Numbers on x-axis represent the variation proportion among detected strains.

### Global carbohydrate utilization potentials were not correlated with member abundances whilst AX-related hydrolytic potentials were broadly dispersed in key member MAGs

CAZymes, especially glycoside hydrolases (GHs), were annotated in each MAG to determine whether member fitness was associated with their genetic potential for carbohydrate degradation. RSAX MAGs generally contained higher abundances of corresponding CAZyme families than WSAX MAGs although all queried GH families were discovered in both metagenomes in a similar distribution pattern (Fig. 5A), suggesting that subtle structural variance did not select communities with broadly distinct carbohydrate utilization capacities. Interestingly, high numbers of CAZyme genes in an organism were not linked with high abundance (indicated by read coverage) in communities (Fig. S6). *Bacteroides* spp. exhibited the greatest diversity of carbohydrate utilization potential, but this did not translate into large populations; *B. ovatus* and *B. thetaiotaomicron* MAGs each contained > 300 CAZyme genes in their MAGs across all metagenomes, but their community abundances were much lower than *A. rectalis* and some bifidobacteria, especially in WSAX consortia.

**Figure 5:**
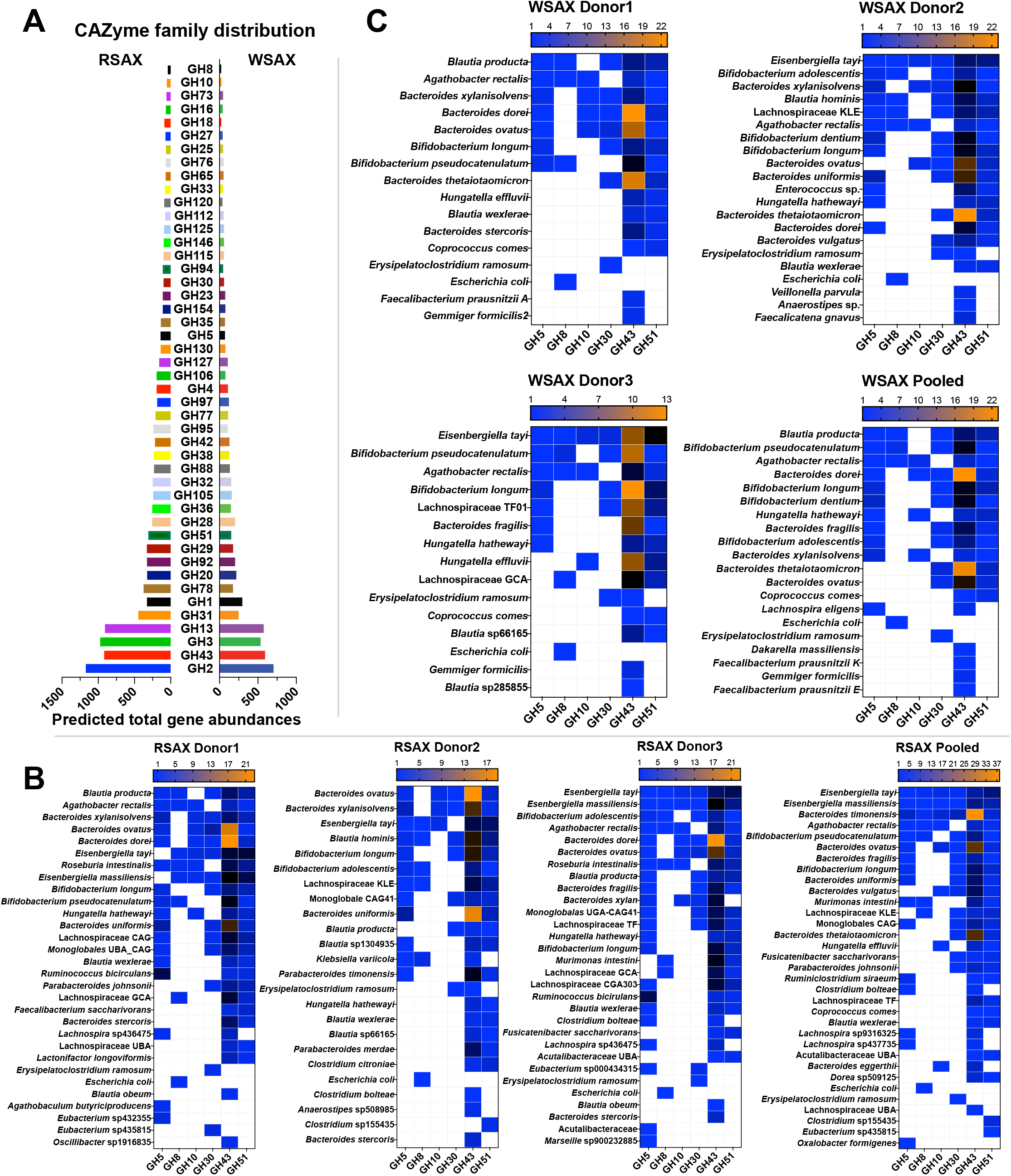
Total predicted CAZyme hydrolase genes among all metagenomic assemblies (A), putative AX backbone degradation CAZymes include glycoside hydrolase (GH) family 10, 8 and 5, in which GH10 is specific to endo-β-1,4-xylanase function, whereas side-chain and terminal xylose hydrolysis genes are in family GH43, GH51. Distribution of six AX-related CAZyme genes in reconstructed MAGs across donors for RSAX (B) and WSAX (C). CAZyme gene numbers were colored from low to high with blue to yellow shades. White cells mean zero detection of such CAZyme gene in genomes. Members not listed in figures were not detected with any of these AX hydrolytic genes.

Not all CAZyme families are likely involved in AX consumption; keystone hydrolytic enzymes with high prevalence in consortial metagenomes include endo-xylanases (e.g., GH5, GH8, GH10 and GH30) and side-chain hydrolases (e.g., GH43 and GH51), which can cleave different glycosidic linkages of AX to produce bacterially accessible oligomers [14, 15]. These AX degradation genes were repeatedly found in MAGs of several shared genera among different consortia (Fig. 5B&C), namely *Agathobacter, Eisenbergiella, Bacteroides, Blautia* and *Bifidobacterium*. MAGs with the same predicted taxonomy across different donors showed similar AX-utilization gene distributions, suggesting that, although they may represent different strains, their core genome potential to consume AX is likely very similar.

GH10-family genes were limited to MAGs of *A. rectalis, E. tayi*, and several *Bacteroides* spp. in WSAX communities, whereas RSAX consortia contained more MAGs with these genes, such as *Roseburia, Hungatella* and different species of *Bacteroides*. This result is possibly due to variant fiber structural configurations in RSAX, which require a greater diversity of xylan backbone-hydrolyzing enzymes. However, genomes of *A. rectalis* did not contain more copies of GH10-family genes than competitors, despite their high abundances. Interestingly, another putative xylanase gene (GH8) was not found in any *Bacteroides* MAGs but was commonly found in Lachnospiraceae spp. (e.g., *A. rectalis* and *Blautia*) and *Bifidobacterium* spp.; this genetic potential might partially explain the relatively high abundances of such organisms in both RSAX and WSAX consortia and suggest potential keystone functions for these organisms in initiating AX consumption. Notably, a MAG identified as *Escherichia coli* also contained a GH8 gene, suggesting this strain might possess AX hydrolysis capacity to some extent.

In contrast to endoxylanase genes, side-chain hydrolases showed much higher gene copy numbers in MAGs, especially GH43-family genes, which were amongst the most abundant CAZyme in assemblies (Fig. 5A). High abundance of GH43 genes suggests the potential of bearing organisms to access more diverse branch linkages in SAX. MAGs from genus *Bacteroides*, including *B. dorei, B. ovatus*, and *B. uniformis*, generally contained relatively higher numbers of GH43 genes (≥ 10) among all MAGs, while *B. fragilis* MAGs contained the fewest GH43 genes amongst *Bacteroides*. Diverse MAGs from genus *Bacteroides* with high GH43 counts were more commonly reconstructed from RSAX consortia compared to the paired WSAX ones. Together, these data suggested 1) that the organisms with the largest repertoires of putative complex SAX-degrading genes were not necessarily the most fit under continuous selection, and 2) that members with large GH43 repertoires may partition niches via degenerate gene functions in ways that relate to SAX structure or via differential gene regulation.

### Domain structures of GH10 and GH43 genes varied across members

CAZyme family classification is not sufficient to fully elucidate polysaccharide utilization potentials of consortia from the two SAXs, since common categories of GH families/sub-families are defined by their core GH domain sequences [16]. Adjacent protein domains fused to GH catabolic domains are also anticipated to influence overall enzyme function, for example in substrate binding and/or hydrolysis. To determine whether accessory domains neighboring GH motifs might differentiate organismal function, we aligned and clustered unique GH10 and GH43 sequences from metagenomes and examined their fused protein domains.

Clustering of GH10 revealed five distant clades, two of which were further divided into phylogenetically constrained subclades (Fig. 6A), suggesting AX backbone hydrolysis functions were largely vertically inherited. Clades were defined by difference in overall domain structure. Genes in clade “L3” and “B1” had different arrangements of carbohydrate-binding modules (CBM). The clade “B1,” found in *Bacteroides* possessed one or more central CBMs between two distinct GH10 domains, an arrangement which might favor a particular AX backbone structure that *Bacteroides* spp. are specialized to degrade. In addition, clade “B2” contained different hydrolase domains besides GH10, such as GH43 and carbohydrate esterase, perhaps targeting glycosidic hydrolysis function towards specific polysaccharide structures (e.g., substituted backbones and branch linkages with esterification). Interestingly, although GH10 genes in each clade were found in both RSAX- and WSAX-degrading cultures, many of them were exclusive to RSAX-degrading cultures. A number of these may reflect the reduced diversity of *Bacteroides* spp. in WSAX consortia, whereas a number of Lachnospiraceae GH10 genes were also only reconstructed from RSAX consortia, likely owing to RSAX’s greater linkage complexity. GH10-containing proteins with predicted signal sequences for secretion were overwhelmingly associated with members of *Bacteroides*.

**Figure 6:**
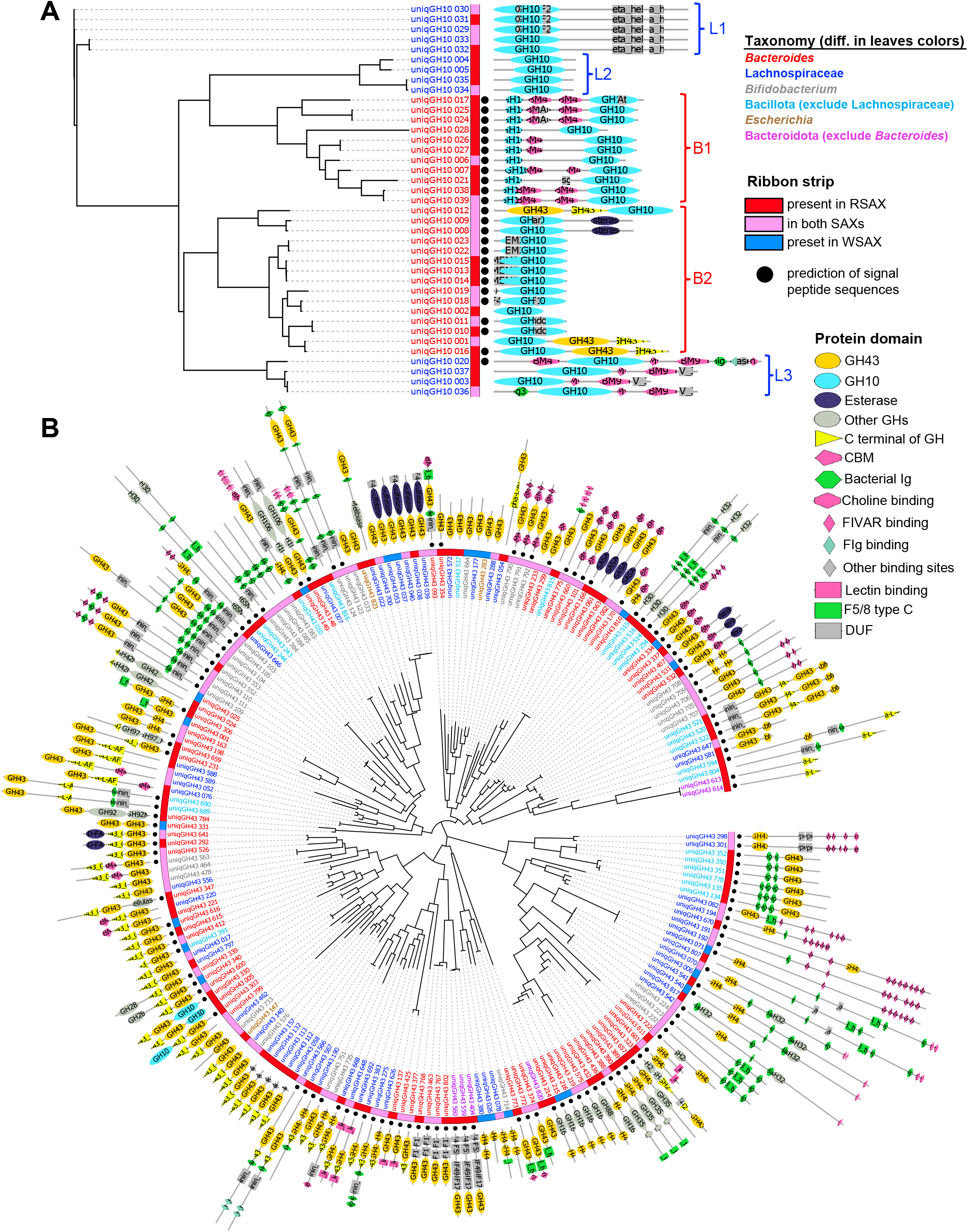
Phylogenetic trees (maximum-likelihood method) of GH10 (A) and GH43 (B) from unique gene protein sequences in day-7 metagenomes. Total 39 GH10 genes and 823 GH43 sequences were collected from all SAX-fermenting metagenomes. The tree of GH43 here represented clustering among unique domain structures, whereas details of gene clusters within simple domain structures were exhibited in Fig. S7. Labels of each leaf are colored at different taxonomy levels: *Bacteroides* (red), Lachnospiraceae (blue), *Bifidobacteria* (grey), Bacillota except Lachnospiraceae(cyan), *Escherichia* (brown) and Bacteroidota exclude *Bacteroides* (magenta). Strips wrapped on leaves display substrate-related information. Black dots are prediction of presence of bacterial signal peptide sequences. Pfam-scanned protein domain structures are illustrated for each leaf.

Multiple different domain architectures of GH43-containing enzymes were observed among AX-degrading consortia, with a total of 88 distinct domain structures from diverse bacterial taxa (Fig. 6B & Fig. S7). Unlike GH10, we observed some GH43 genes specific to either RSAX- or WSAX-consuming consortia, as well as some shared by both. Genes encoding predicted secretion signals were found across phyla. Genes with simpler module structures (Fig. S7A) were grouped together across taxa, whilst genes with multiple fused domains were more taxon-specific (Fig. S7B). The GH43 domain-only structures occupied a variety of positions within their overall open reading frames. For example, in a cluster of genes containing dual GH43 domain fusions (Fig. S7H), a clade with shorter GH43 domains was dominated by MAGs from Lachnospiraceae and *Bifidobacterium*, while the other clades were composed of *Bacteroides*. Interestingly, this cluster also showed a substrate-dependent grouping effect, in which the Lachnospiraceae and bifidobacteria clade was found in WSAX-fermenting consortia. GH43|CBM6 fusions were entirely dominated by members of *Bacteroides*, as were GH43|F5/8 typeC fusions (the latter with one exception from *Parabacteroides*). With respect to GH43 genes with multiple domain structures (Fig. S7B), majority module configurations were conserved within taxa, whilst few similar motif structures were observed across taxa (Fig. 5B). For example, a protein structure of CBM4|GH43|CBM6|FIVAR (from *Bifidobacterium*) was clustered closely with CBM6|GH43|CBM6|CBM6 (from *Bacteroides*), both containing a GH43 domain in-between substrate binding modules, suggesting possible convergent evolution. Taken together, these results suggested that GH-containing genes are not functionally equivalent and may be specific to very fine polysaccharide structural differences.

## DISCUSSION

Improved understanding of how complex dietary fibers impact gut community assembly and members’ DOL is essential to decipher the ecological mechanisms driving inter-individual differences in response to fibers and to engineer fibers for predictable, beneficial microbiome outcomes at population scales. Here, we attempted to determine to what extent fiber structures governed community assembly across individual microbiomes using sequential *in vitro* batch fermentation. This approach has been proven to select substrate-specific consortia derived from the human gut microbiome; our previous study revealed stable and high diversity SAX-fermenting communities using this approach, in contrast to low-diversity communities cultured on simpler substrates [7]. One limitation of this proof-of-concept study, however, was its examination of a single AX polysaccharide and a single donor’s fecal inoculum. Human gut microbiota vary widely among healthy individuals [17], and ecological founder effects have been proposed to drive much of the personalized fiber responses that have been widely reported in human studies [18, 19]. Although a structurally complex dietary fiber is anticipated to initiate shifts in microbial community structure [20] based upon which organisms possess the genetic potential to degrade it in part or whole, the extent to which fine fiber structural variants diverge in their impact is poorly understood. We report that similar polysaccharides with even subtle structure variances can differentially modulate the membership and metabolic function of microbial consortia. Notably, communities selected using the same substrate converge in membership across individuals, despite divergent initial starting points (Fig. S3A). This finding advances the hypothesis that a shared human core gut microbiota exists across diverse individuals, which is populated by identical or nearly-identical strains (although some population sizes in initial fecal samples are below the level of detection by sequencing-based approaches), despite known strain-level competition in the gut [21, 22]. If true, it would suggest that fine structural variants of complex fibers may, over time, be used to target consortia, or even specific species or strains, across individuals. Two possible mechanisms for the cross-donor convergence are that 1) AX structures drive initial selection of microbes with cognate CAZyme genes as keystone species, and 2) such selection orders a diverse cross-feeding community with metabolic exchanges based upon the outputs of these primary organisms (e.g., partly degraded AX-oligosaccharides, variant organic acids as electron sinks, and other nutrients). Similar role division into primary degrader, secondary degrader, and generalist species has been observed in degradation of structurally less-complex polysaccharides in particles by marine microbes [10, 23–25]. Our systems may exhibit increased DOL to this system in that 1) lack of obvious terminal electron acceptors for respiration (besides CO_2_ and trace SO_4_^2-^) means that exchange of intermediately reduced organic molecules as electron sinks may foster mutualism, 2) densely branched polysaccharides may require sequential enzymatic activity for degradation, which may reside across members, and 3) the increased structural diversity of the oligosaccharide pool may permit more diversity in secondary degraders. However, the lack of the same ability to spatially organize on soluble polysaccharides may drive comparative reductions in DOL in our system. In general, however, our study suggests very similar membership – down to the strain level – across three independent donors, suggesting fewer generalist roles (e.g. oxidation of organic acids metabolized for energy [24]) may exist in our system.

Further, our study demonstrates that divergent fiber structures select for organisms bearing CAZymes with varying domain structure, suggesting that these enzymes are not functionally redundant in their interactions with polysaccharides. Rather, their domain structures may differentially bind certain motifs and cleave around side chains that might otherwise exert steric hindrance. Interactions between the keystone species and fiber structures may then determine the substrates available to cross-feed other members and metabolic exchanges within communities. As an example, endoxylanase-producing species, such as *A. rectalis* and *Bacteroides* spp., may differentially initiate AX degradation, in the process generating a distinct suite of smaller oligosaccharides available to non-endoxylanase producers [26]. These secondary consumers (e.g. *B. longum*) may, in turn, support the growth of endoxylanase producers by providing electron acceptors they can further reduce, such as lactate (which can be further converted either to butyrate or propionate) or acetate (which is likely used by *A. rectalis* and other Lachnospiraceae members to produce butyrate) [27]. Similar behavior was reported in *Bacteroides thetaiotaomicron* support of bifidobacteria growing in its spent medium; however, glycan structure appears to dictate whether *Bacteroides* spp. will exhibit cooperative or selfish behavior [28]. Interestingly, large amounts of lactate accumulated in WSAX consortia (Fig. S4), possibly indicating less lactate consumption in WSAX communities or a faster lactate production rate by larger populations of *Bifidobacterium* species; although the hypothetical cross-feeding pair of *A. rectalis* and *Bifidobacterium* spp. were found in both, the abundance of two organisms was much higher in WSAX than RSAX consortia. These cross-feeding interactions may impart stability to communities, especially as members may display functional degeneracy in metabolite exchange even as they exhibit specialization in CAZyme expression. Differential metabolite concentrations (e.g., of lactate and SCFAs) in RSAX and WSAX samples suggest that the secondary metabolites available in each consortium for cross-feeding are likely distinct.

Hydrolytic functions of CAZymes have been widely established with their affinity for different polysaccharide recognition [29–31] at the level of CAZyme families (e.g. GH10, GH43) containing enzymes of similar activity. However, our study reveals that the other domains fused to GH domains vary across taxa and are differentially found in cultures consuming the two different SAX types (Fig. 5A). Thus, our study suggests the hypothesis that fine polysaccharide structure interactions with functionally degenerate, but not redundant, genome-encoded CAZyme complements to impart selective forces on community assembly. In our AX-degrading consortia, endoxylananse (GH10)-encoding species were limited to *Bacteroides* spp. and Lachnospiraceae spp. (mainly *A. rectalis*) with putative distinct substrate-binding motifs (CBM4 of *Bacteroides* spp. vs. CBM9 of *A. rectalis*) (Fig. 6A). This suggests that degradation of the xylan backbone was limited to a few specific taxa with possible structural specificities, leading to distinct oligosaccharide hydrolysate pools that may drive divergent cross-feeding patterns. Distinct GH10 domain structures in AX consortia were likely vertically inherited, from which we further hypothesize that altering substitution (branch) structure on the xylan backbone could potentially result in selection of different dominant backbone-consuming organisms (i.e., *Bacteroides* spp. vs. *A. rectalis*), thereby determining community metabolic function. With respect to GH43 domain motifs, multiple putative substrate binding sites beyond CBM domains were identified between both primary degraders and consumers without endoxylanse function (Fig. S7B). Such non-CBM substrate binding modules include “laminin_G_3”, “FIVAR”, “Big_2”, etc., and may alter the affinity or activity of GH catabolic domains with respect to local structural motifs. For example, “laminin_G_3” selectively binds with 3^2^-α-l-arabinofuranosyl-xylobiose and 2^3^,3^3^-di-α-l-arabinofuranosyl-xylotriose [32], suggesting a potential hydrolytic target of branched arabinofuranosyl residues on xylooligomers. “FIVAR” domains have been extensively reported along with CAZymes and are thought to have carbohydrate binding functions, but are not yet well-characterized [33–35]. “Big_2” is a immunoglobulin(Ig)-like domain that is also widely found in CAZymes of gut origin [36, 37], including amylases, chitinases, xylanases, and cellulases, and this module may behave multifunctionally in aiding the function of GH reactive motifs, such as membrane-anchoring and improvement of thermal-stability between catalytic center and substrate [37, 38]. The availability of diverse potential carbohydrate binding motifs in AX-derived GH43 gene pools indicate a vast and degenerate genomic potential for AX degradation, which may drive specialization and DOL [12]. Highly diverse domain fusions may mediate differences in substrate-utilization function of multiple copies of GH43-containing CAZymes in genomes and, further, similarities in alternate domain content across large phylogenetic distances may suggest which organisms may compete vs. cooperate for specific AX structural motifs. Overall, our study suggests that high dilution pressure, sequential *in vitro* fermentations are a powerful tool to enrich core polysaccharide-specific consortia across individuals’ gut diverse microbiomes and may help to associate different GH43 domain fusions with their structural specificity, over time establishing a more mechanistic model for fine polysaccharide structure impacts on microbiome structure and function. More generally, this approach may permit the selection and engineering of functional polysaccharides (including, but not limited to, diverse hemicelluloses) for predictable gut microbiome impact across idiosyncratic individuals and at population scales.

## METHODS

### Polysaccharide purification and chemical structure determination

Polymer arabinoxylans (AXs) were isolated from two sorghum varieties (P721N-2016 [phenotype “red”] and P850637-2017 [phenotype “white”] Purdue Agronomy Center). Soluble AXs were extracted following an order of de-fat (hexane), enzymatic de-starch and protein removal (amylase and proteinase), alkaline incubation (1M NaOH, 60 °C for 4 h) and with ethanol precipitation as described previously [13]. The alkaline-soluble fraction isolated from red and white sorghum brans were codified as “RSAX” and “WSAX”, respectively, and were stored under −20 °C for future analyses.

Neutral sugar composition of two SAXs were characterized by acid hydrolysis (2M trifluoracetic acid, 121 °C, 90 min) followed by a reduction process and alditol acetate derivatization [39]. Glycosidic linkage profiles of SAXs were determined as partially methylated alditol acetates [39] with four steps (i.e., methylation, hydrolysis, reduction and acetylation). A gas chromatograph-mass spectrometer (GC-MS) equipped with a SP 2330 column and a triple-axis detector (Agilent Technologies, Santa Clara, CA, USA) was used to measure both sugar composition and relative proportions of linkage indicates. Helium was used as carrier gas (1 mL/min). Oven temperature was set at 170 °C for 2 min, followed by a ramp up to 260 °C with 3 °C/min and held for 2 min.

### *In vitro* sequential batch fermentation

An upper GI digestion of substrates was performed before fermentation with amylase, pepsin and pancreatin [40]. Digesta were dialyzed (6-8 KDa cut-off) and freeze-dried. Fecal fermentations were performed in an anaerobic chamber (Coy Laboratory Products Inc., Great Lake, MI) with mixed anaerobic gas (90% N_2_, 5% CO_2_ and 5% H_2_). A chemically defined gut mineral medium was prepared [7], with buffer strength of 40 mM sodium phosphate and 1% AX as sole carbohydrate (complete formulation shown in Supplementary Information).

Human fecal samples were collected fresh from three healthy donors. Donors (ages 20– 25) had not received any antibiotic treatment within three months and had not consumed prebiotic or probiotic products for two weeks. The protocol for collecting human feces were approved by Institutional Review Board of Purdue University (#1906022302). Fresh fecal samples were sealed and transferred into anaerobic chamber immediately and used within 30 min. Fecal inocula were prepared by diluting feces with media (1:20, w/v) and filtered with cheesecloth. A pooled inoculum was prepared by evenly mixing inocula from three donors. Fecal inocula (1 mL) were further mixed with 4 mL of medium in tubes to obtain a final dilution rate of 1:100 (w/v).

*In vitro* consecutive batch fermentation was carried out in sterilized Balch tubes as previously reported [7]. Briefly, anaerobic tubes were incubated at 37 °C (150 rpm) for 24 h, and then passaged to fresh media at a 1:100 dilution rate every 24 h, continued for a total of 7 passages (Fig 1C). Each sample was cultivated in triplicate lineages that were not intermixed.

### DNA extraction and 16S amplicon sequencing

Consortial DNA was extracted using a phenol-chloroform method as in previous studies [41, 42] with minor modifications [7] (detailed protocol in supplementary information). Briefly, cell pellets were enzymatically lysed, followed by a 10 sec bead beating step before phenol-chloroform purification. Purified DNA fraction was alcohol precipitated and then stored in Tris-EDTA buffer at −20 °C.

Library preparation of 16S rRNA gene amplicons was performed as described previously [43]. Two primers 515-FB (GTGYCAGCMGCCGCGGTAA) and 926-R (CCGYCAATTYMTTTRAGTTT) were used to amplify the V4-V5 region [44]. Resulting amplicons were purified by an AxyPrep PCR Clean-up Kit (Axygen Inc., Tewksbury, MA). Amplicons were then barcoded using the TruSeq dual-index approach and purified as above. Barcoded amplicons were quantified using a Qubit dsDNA HS Assay Kit (Invitrogen, Carlsbad, CA) and pooled accordingly. Pool quality was ascertained using an Agilent Bioanalyzer (Agilent, Santa Clara, CA) and sequencing was performed on an Illumina MiSeq platform (2 × 250 cycles) at the Purdue Genomics Core Facility. Amplicon sequences were processed by mothur v.1.39.3 [45] following the MiSeq SOP with the modifications described previously [7] (full analysis protocol in supplementary information).

### Shotgun metagenomics and genome reconstruction

Shotgun sequencing libraries were prepared using the Illumina TruSeq DNA Nano kit and then sequenced on an Illumina NovaSeq 6000 S4 platform (2 × 150 bp). Sequencing processes of *de novo* contigs assembly, genome reconstruction and manual refinement, and functional annotation are detailed in Supplementary Information. Briefly, raw reads were assembled using metaSPAdes (3.14.1) and statistics calculated using QUAST (3.2) [46]. Metagenome-assembled genomes (MAGs) were reconstructed with MaxBin2 (2.2.3) [47] and refined with RefineM (0.1.2) [48]. Gene models were predicted by Prodigal (2.6.3) [49] and genes were annotated by HMMer (3.1b2) [50] using Pfam (33.1) [51] and TIGRFAM (15.0) [52] databases. MAG quality was visualized and manually refined using Anvi’o (ver. 6.2) [53] (MAGs information in Supplementary Table S1). Strain-level relationships were determined using StrainPhlAn (3.0) [54]. CAZyme gene annotation was performed using HMMer with the dbCAN2 database (V8) [55]. Phylogenetic trees were built from protein sequence alignment of CAZyme genes (Muscle-v5) [56] with maximum-likelihood joining using FastTree (2.1.11) [57].

### Community metabolites and non-targeted metabolomics

The concentration of acetate, propionate and butyrate in the fermentations was measured as described previously [43]. Samples were measured on a GC (Agilent 7890A) with a fused silica column (Supelco 40369-03A).

Non-targeted metabolomics were measured on a Thermo Trace 1310 GC (Thermo Fisher Scientific Inc., Waltham, MA) coupled with Thermo TSQ 8000 triple quadrupole mass spectrometer (Thermo Fisher Scientific Inc., Waltham, MA) at the Purdue Metabolite Profiling Facility. Briefly, samples were subjected to Bligh-Dyer extraction, followed with a trimethylsilyl (TMS) derivatization, and then measured immediately on GC-MS (details in Supplementary Information). Cluster on metabolomics over groups were built using R 4.1.0 with Euclidian distance.

## Acknowledgements

The authors gratefully acknowledge grant support from the National Institute of General Medical Sciences through an Early Career Maximizing Investigator’s Research Award (R35 GM133634) to SRL, the U.S. Department of Agriculture through Hatch project IND011670 to SRL, and institutional funds provided by the Purdue University Departments of Food Science and Nutrition Science. TY is grateful for support from the CSC (China Scholarship Council) program. The authors are thankful to Laura Libera for assistance with arabinoxylan extraction, fermentation experiments, and sample collection.

## Data Availability Statement

All raw sequencing data in FASTQ format were deposited in the NCBI Sequence Read Archive (SRA) under BioProject PRJNA866162 as BioSamples SRX16850208-SRX16849926.

## Authorship contribution statement

Tianming Yao: Investigation, Methodology, Conceptualization, Data collection, Data curation, Formal analysis, Writing - original draft, Writing - editing. Dane G. Deemer: Data visualization, Formal analysis, Writing - reviewing. Ming-Hsu Chen: Investigation, Methodology, Conceptualization, Data collection, Writing - reviewing. Bradly L. Reuhs: Methodology, Data curation, Writing – reviewing. Bruce R. Hamaker: Methodology, Data curation, Writing – reviewing. Stephen R. Lindemann: Project administration, Supervision, Conceptualization, Investigation, Methodology, Writing - original draft, Writing - review & editing.

## Notes

### Competing Interest Statement

Dr. Lindemann was been funded by the NIH. He also sits on the scientific advisory boards of the Grain Foods Foundation, CP Kelco, US and OLIPOP Inc. and has received compensation in those roles, and founded and owns Ina Microbiome LLC, a microbiome consulting firm. Dr. Hamaker sits on the scientific advisory board of the Nebraska Food for Health Center and is a partner in RightCarbs LLC. Mr. Deemer and Drs. Yao, Chen, and Reuhs declare no potential competing interests. The funders played no role in the design or execution of the study or the decision to publish.

## REFERENCE

1. Huttenhower C, Gevers D, Knight R, Abubucker S, Badger JH, Chinwalla AT, et al. Structure, function and diversity of the healthy human microbiome. Nature 2012; 486: 207–214.

2. Kamada N, Chen GY, Inohara N, Núñez G. Control of pathogens and pathobionts by the gut microbiota. Nat Immunol 2013; 14: 685–690.

3. Sivaprakasam S, Prasad PD, Singh N. Benefits of short-chain fatty acids and their receptors in inflammation and carcinogenesis. Pharmacol Ther 2016; 164: 144–151.

4. Tian Y, Xu Q, Sun L, Ye Y, Ji G. Short-chain fatty acids administration is protective in colitis-associated colorectal cancer development. J Nutr Biochem 2018; 57: 103–109.

5. Brooks L, Viardot A, Tsakmaki A, Stolarczyk E, Howard JK, Cani PD, et al. Fermentable carbohydrate stimulates FFAR2-dependent colonic PYY cell expansion to increase satiety. Mol Metab 2017; 6: 48–60.

6. Kim CH. Microbiota or short-chain fatty acids: which regulates diabetes? Cell Mol Immunol 2018; 15: 88–91.

7. Yao T, Chen M-H, Lindemann SR. Structurally complex carbohydrates maintain diversity in gut-derived microbial consortia under high dilution pressure. FEMS Microbiol Ecol 2020; 96.

8. Cantu-Jungles TM, Bulut N, Chambry E, Ruthes A, Iacomini M, Keshavarzian A, et al. Dietary Fiber Hierarchical Specificity: the Missing Link for Predictable and Strong Shifts in Gut Bacterial Communities. mBio; 12: e01028–21.

9. Romero Marcia AD, Yao T, Chen M-H, Oles RE, Lindemann SR. Fine Carbohydrate Structure of Dietary Resistant Glucans Governs the Structure and Function of Human Gut Microbiota. Nutrients 2021; 13: 2924.

10. Pontrelli S, Szabo R, Pollak S, Schwartzman J, Ledezma-Tejeida D, Cordero OX, et al. Metabolic cross-feeding structures the assembly of polysaccharide degrading communities. Sci Adv 2022; 8: eabk3076.

11. Abbott DW, van Bueren AL. Using structure to inform carbohydrate binding module function. Curr Opin Struct Biol 2014; 28: 32–40.

12. Lindemann SR. A piece of the pie: engineering microbiomes by exploiting division of labor in complex polysaccharide consumption. Curr Opin Chem Eng 2020; 30: 96–102.

13. Rumpagaporn P, Reuhs BL, Kaur A, Patterson JA, Keshavarzian A, Hamaker BR. Structural features of soluble cereal arabinoxylan fibers associated with a slow rate of in vitro fermentation by human fecal microbiota. Carbohydr Polym 2015; 130: 191–197.

14. Lagaert S, Pollet A, Delcour JA, Lavigne R, Courtin CM, Volckaert G. Substrate specificity of three recombinant α-l-arabinofuranosidases from Bifidobacterium adolescentis and their divergent action on arabinoxylan and arabinoxylan oligosaccharides. Biochem Biophys Res Commun 2010; 402: 644–650.

15. Lagaert S, Pollet A, Courtin CM, Volckaert G. β-Xylosidases and α-l-arabinofuranosidases: Accessory enzymes for arabinoxylan degradation. Biotechnol Adv 2014; 32: 316–332.

16. Aspeborg H, Coutinho PM, Wang Y, Brumer H, Henrissat B. Evolution, substrate specificity and subfamily classification of glycoside hydrolase family 5 (GH5). BMC Evol Biol 2012; 12: 186.

17. Arumugam M, Raes J, Pelletier E, Paslier DL, Yamada T, Mende DR, et al. Enterotypes of the human gut microbiome. Nature 2011; 473: 174–180.

18. Salonen A, Lahti L, Salojärvi J, Holtrop G, Korpela K, Duncan SH, et al. Impact of diet and individual variation on intestinal microbiota composition and fermentation products in obese men. ISME J 2014; 8: 2218–2230.

19. Fransen F, van Beek AA, Borghuis T, Meijer B, Hugenholtz F, van der Gaast-de Jongh C, et al. The Impact of Gut Microbiota on Gender-Specific Differences in Immunity. Front Immunol 2017; 8.

20. Cantu-Jungles TM, Hamaker BR. New View on Dietary Fiber Selection for Predictable Shifts in Gut Microbiota. mBio 2020; 11.

21. Walter J, Maldonado-Gómez MX, Martínez I. To engraft or not to engraft: an ecological framework for gut microbiome modulation with live microbes. Curr Opin Biotechnol 2018; 49: 129–139.

22. Garud NR, Good BH, Hallatschek O, Pollard KS. Evolutionary dynamics of bacteria in the gut microbiome within and across hosts. PLOS Biol 2019; 17: e3000102.

23. Sichert A, Cordero OX. Polysaccharide-Bacteria Interactions From the Lens of Evolutionary Ecology. Front Microbiol 2021; 12: 705082.

24. Enke TN, Datta MS, Schwartzman J, Cermak N, Schmitz D, Barrere J, et al. Modular Assembly of Polysaccharide-Degrading Marine Microbial Communities. Curr Biol 2019; 29: 1528-1535.e6.

25. Ebrahimi A, Schwartzman J, Cordero OX. Cooperation and spatial self-organization determine rate and efficiency of particulate organic matter degradation in marine bacteria. Proc Natl Acad Sci U S A 2019; 116: 23309–23316.

26. Rivière A, Gagnon M, Weckx S, Roy D, Vuyst LD. Mutual Cross-Feeding Interactions between Bifidobacterium longum subsp. longum NCC2705 and Eubacterium rectale ATCC 33656 Explain the Bifidogenic and Butyrogenic Effects of Arabinoxylan Oligosaccharides. Appl Environ Microbiol 2015; 81: 7767–7781.

27. Fukuda S, Toh H, Hase K, Oshima K, Nakanishi Y, Yoshimura K, et al. Bifidobacteria can protect from enteropathogenic infection through production of acetate. Nature 2011; 469: 543–547.

28. Luis AS, Briggs J, Zhang X, Farnell B, Ndeh D, Labourel A, et al. Dietary pectic glycans are degraded by coordinated enzyme pathways in human colonic Bacteroides. Nat Microbiol 2018; 3: 210.

29. Sørensen HR, Jørgensen CT, Hansen CH, Jørgensen CI, Pedersen S, Meyer AS. A novel GH43 α-l-arabinofuranosidase from Humicola insolens: mode of action and synergy with GH51 α-l-arabinofuranosidases on wheat arabinoxylan. Appl Microbiol Biotechnol 2006; 73: 850–861.

30. Mechelke M, Koeck DE, Broeker J, Roessler B, Krabichler F, Schwarz WH, et al. Characterization of the arabinoxylan-degrading machinery of the thermophilic bacterium Herbinix hemicellulosilytica—Six new xylanases, three arabinofuranosidases and one xylosidase. J Biotechnol 2017; 257: 122–130.

31. Holck J, Djajadi DT, Brask J, Pilgaard B, Krogh KBRM, Meyer AS, et al. Novel xylanolytic triple domain enzyme targeted at feruloylated arabinoxylan degradation. Enzyme Microb Technol 2019; 129: 109353.

32. Sakka M, Kunitake E, Kimura T, Sakka K. Function of a laminin_G_3 module as a carbohydrate-binding module in an arabinofuranosidase from Ruminiclostridium josui. FEBS Lett 2019; 593: 42–51.

33. Chitayat S, Adams JJ, Furness HS, Bayer EA, Smith SP. The solution structure of the C-terminal modular pair from Clostridium perfringens μ-toxin reveals a noncellulosomal dockerin module. J Mol Biol 2008; 381: 1202–1212.

34. Fujita K, Oura F, Nagamine N, Katayama T, Hiratake J, Sakata K, et al. Identification and molecular cloning of a novel glycoside hydrolase family of core 1 type O-glycan-specific endo-α-N-acetylgalactosaminidase from Bifidobacterium longum. J Biol Chem 2005; 280: 37415–37422.

35. Gaytán MO, Singh AK, Woodiga SA, Patel SA, An S-S, León AV-P de, et al. A novel sialic acid-binding adhesin present in multiple species contributes to the pathogenesis of Infective endocarditis. PLOS Pathog 2021; 17: e1009222.

36. Han Q, Liu N, Robinson H, Cao L, Qian C, Wang Q, et al. Biochemical characterization and crystal structure of a GH10 xylanase from termite gut bacteria reveal a novel structural feature and significance of its bacterial Ig-like domain. Biotechnol Bioeng 2013; 110: 3093–3103.

37. Xing M, Wang Y, Zhao Y, Chi Z, Chi Z, Liu G. C-Terminal Bacterial Immunoglobulin-like Domain of κ-Carrageenase Serves as a Multifunctional Module to Promote κ-Carrageenan Hydrolysis. J Agric Food Chem 2022; 70: 1212–1222.

38. Wakinaka T, Kiyohara M, Kurihara S, Hirata A, Chaiwangsri T, Ohnuma T, et al. Bifidobacterial α-galactosidase with unique carbohydrate-binding module specifically acts on blood group B antigen. Glycobiology 2013; 23: 232–240.

39. Pettolino FA, Walsh C, Fincher GB, Bacic A. Determining the polysaccharide composition of plant cell walls. Nat Protoc 2012; 7: 1590.

40. Rose DJ, Patterson JA, Hamaker BR. Structural differences among alkali-soluble arabinoxylans from maize (Zea mays), rice (Oryza sativa), and wheat (Triticum aestivum) brans influence human fecal fermentation profiles. J Agric Food Chem 2009; 58: 493–499.

41. Ferrera I, Massana R, Balagué V, Pedrós-Alió C, Sánchez O, Mas J. Evaluation of DNA extraction methods from complex phototrophic biofilms. Biofouling 2010; 26: 349–357.

42. Lindemann SR, Moran JJ, Stegen JC, Renslow RS, Hutchison JR, Cole JK, et al. The epsomitic phototrophic microbial mat of Hot Lake, Washington: community structural responses to seasonal cycling. Front Microbiol 2013; 4: 323.

43. Thakkar RD, Tuncil YE, Hamaker BR, Lindemann SR. Maize Bran Particle Size Governs the Community Composition and Metabolic Output of Human Gut Microbiota in in vitro Fermentations. Front Microbiol 2020; 11.

44. Walters W, Hyde ER, Berg-Lyons D, Ackermann G, Humphrey G, Parada A, et al. Improved bacterial 16S rRNA gene (V4 and V4-5) and fungal internal transcribed spacer marker gene primers for microbial community surveys. Msystems 2016; 1: e00009–15.

45. Schloss PD, Westcott SL, Ryabin T, Hall JR, Hartmann M, Hollister EB, et al. Introducing mothur: open-source, platform-independent, community-supported software for describing and comparing microbial communities. Appl Environ Microbiol 2009; 75: 7537–7541.

46. Gurevich A, Saveliev V, Vyahhi N, Tesler G. QUAST: quality assessment tool for genome assemblies. Bioinforma Oxf Engl 2013; 29: 1072–1075.

47. Wu Y-W, Simmons BA, Singer SW. MaxBin 2.0: an automated binning algorithm to recover genomes from multiple metagenomic datasets. Bioinformatics 2016; 32: 605–607.

48. Parks DH, Rinke C, Chuvochina M, Chaumeil P-A, Woodcroft BJ, Evans PN, et al. Recovery of nearly 8,000 metagenome-assembled genomes substantially expands the tree of life. Nat Microbiol 2017; 2: 1533–1542.

49. Hyatt D, Chen G-L, LoCascio PF, Land ML, Larimer FW, Hauser LJ. Prodigal: prokaryotic gene recognition and translation initiation site identification. BMC Bioinformatics 2010; 11: 119.

50. Johnson LS, Eddy SR, Portugaly E. Hidden Markov model speed heuristic and iterative HMM search procedure. BMC Bioinformatics 2010; 11: 431.

51. Finn RD, Bateman A, Clements J, Coggill P, Eberhardt RY, Eddy SR, et al. Pfam: the protein families database. Nucleic Acids Res 2014; 42: D222–D230.

52. Haft DH, Selengut JD, White O. The TIGRFAMs database of protein families. Nucleic Acids Res 2003; 31: 371–373.

53. Eren AM, Esen ÖC, Quince C, Vineis JH, Morrison HG, Sogin ML, et al. Anvi’o: an advanced analysis and visualization platform for ‘omics data. PeerJ 2015; 3: e1319.

54. Beghini F, McIver LJ, Blanco-Míguez A, Dubois L, Asnicar F, Maharjan S, et al. Integrating taxonomic, functional, and strain-level profiling of diverse microbial communities with bioBakery 3. eLife 2021; 10: e65088.

55. Zhang H, Yohe T, Huang L, Entwistle S, Wu P, Yang Z, et al. dbCAN2: a meta server for automated carbohydrate-active enzyme annotation. Nucleic Acids Res 2018; 46: W95–W101.

56. Edgar RC. MUSCLE: multiple sequence alignment with high accuracy and high throughput. Nucleic Acids Res 2004; 32: 1792–1797.

57. Price MN, Dehal PS, Arkin AP. FastTree: Computing Large Minimum Evolution Trees with Profiles instead of a Distance Matrix. Mol Biol Evol 2009; 26: 1641–1650.

