## Supplementary methods for "SUBTLE DIFFERENCES IN FINE POLYSACCHARIDE STRUCTURES GOVERN SELECTION AND SUCCESSION OF HUMAN GUT MICROBIOTA"

**SUPPLEMENTARY INFORMATION for METHODS**

**Gut mineral media recipe:**

Base medium (1 L) included 0.47 g NaCl, 0.45 g KCl, 0.10 g Na_2_SO_4_, 0.001 g resazurin, 3.59 g Na_2_HPO_4_ and 1.87 g NaH_2_PO_4_. Both urea (0.40 g) and NH_4_Cl (0.36 g) were added as nitrogen sources, and the pH was adjusted to 7.0 ± 0.1 by NaOH. The base medium was then autoclaved at 121 °C for 20 min. Heat sensitive nutrients, including 20 essential amino acids (200 μM total; 10 μM of each), 1 % (v/v) ATCC vitamin supplement (ATCC MD-VS; Hampton, NH), 0.0728 g CaCl_2_, 0.1 g MgCl_2_, 0.25 g L-cysteine hydrochloride and 1000X P1 trace metal solution (1 mL), were sterilized by 0.22 μm filter and added. P1 metal solution contains (per liter) 34.26 g H_3_BO_3_, 4.32 g MnCl_2_⋅4H_2_O, 315 mg ZnCl_2_, 44 mg Na_2_MoO_4_⋅2H_2_O, 3 mg CuSO_4_⋅5H_2_O, 12.15 mg CoCl_2_⋅6H_2_O, 259 mg NiCl_2_, 0.28 ml EDTA (10 mM), 1 ml FeCl3⋅6H2O (3.89 mg/ml in 0.1 M HCl). Upper GI digested SAXs were added as the sole carbohydrates into media at a concentration of 1 % (w/v).

**Phenol-chloroform DNA extraction method**:

Cell pellets from each sample were incubated at 85°C for 5 min. After cooling, 500 µL lysozyme solution (1 mg/mL) was added into samples and incubated at 37°C for 45 min. A second incubation was followed with 100 µL Proteinase K (1.2 mg/mL) at 56°C for 1 h. The slurries were then combined with 500 µL phenol/chloroform/isoamyl alcohol solution (25:24:1, PCI) in lysis tubes (330 TX, BioSpec Products, Inc., Bartlesville, OK) preloaded with 0.3 g of 0.1 mm silica beads (1107910z, BioSpec Products, Inc., Bartlesville, OK). The lysis tubes were subjected to bead beating in a FastPrep-24 homogenizer (MP Biomedicals, Santa Ana, CA) for 10 sec (6 m/s). An additional 500 µL of PCI solution was added and vortexed for 15 sec. Samples were centrifuged (13,000 × g, 10 min, 4 °C), and the upper aqueous phase was transferred to new tubes and mixed with 1 mL of a chloroform/isoamyl alcohol solution (24:1) by vortex for 15 sec. Samples were centrifuged again (13,000 × g, 10 min) and the upper aqueous phase was transferred to new tubes, 100 µL of 3 M sodium acetate was added, and precipitated with 1 mL of isopropanol. The solution was shaken gently, incubated at room temperature for 10 min, and then centrifuged at 13,000 × g for 30 min. Supernatants were decanted and the DNA pellets were washed with 500 µL 70% ethanol twice. The microbial DNA was then air-dried and resuspended in Tris-EDTA buffer (pH 8.0) and stored at -20 °C.

**16S sequencing reads processing:**

Amplicon reads were processed using mothur v.1.39.3 according to the MiSeq SOP (<https://www.mothur.org/wiki/MiSeq_SOP>) with previously described modifications [1]. Briefly, contigs were built on merging read pairs and errors were screened. Error contigs were discarded with any error detected in primer region, along with any ambiguous bases, homopolymer lengths over 8 bases, and total contig lengths larger than 411 bp. Filtered reads were aligned to SILVA bacteria reference database (v.132, mothur-formatted) [2]. Identical or duplicated aligned reads were merged and unique reads were then pre-clustered with up to 2 differences among unique reads (~99.5% identity). Chimera reads were then searched by mothur-embedded UCHIME algorithm. Checked and filtered reads were further classified using the Ribosomal Database Project (RDP) classifier (version 18, mothur-formatted training set with species epithets) [3, 4]. Taxonomy classifications were cut off at a bootstrap value of 95. Sequences classified to non-bacterial taxa were removed and the final specie-level operational taxonomic unit (OTU) table (shared file) was created for downstream analyses.

Ecological indices were calculated using mothur-embedded algorithms: α-diversity via sobs, chao, simpsoneven, shannon and invsimpson calculators and β-diversity using Bray-Curtis dissimilarity. The group distances and consortia evolution routes based on β-diversity were visualized as bi-plots of non-metric multi-dimensional scaling (NMDS), principal coordinate analysis (PCoA) and redundancy analysis (RDA) using R 4.1.0 (R Foundation for Statistical Computing, Vienna, Austria) with vegan packages.

**Metagenomic contigs assembly, genome reconstruction and functional annotation**

Sequencing reads quality were examined with FastQC (0.11.9), generating 2.216 × 10^9^ quality-filtered reads with total 3.105 × 10^11^ bp. A *de novo* assembly-based approach was used with each sample assembled separately. Raw reads were assembled using metaSPAdes, ver. 3.14.1 with default *k*-mer sizes (21, 33, 55). The assembly statistics for the 8 contig sets were calculated by QUAST v.3.2 [5]. N_50_ length was 53465, 70120, 49098, 77850, 48598, 34692, 31891 and 39254 bp for W1, W2, W3, W-pooled, R1, R2, R3 and R-pooled sample, respectively.

Draft microbial genome reconstructions (metagenome-assembled genomes, MAGs) were generated with MaxBin2 (ver. 2.2.3) [6] using coverages calculated from reads mapping the assembly at 99% identity, and refined with RefineM (ver. 0.1.2) to automatically improve binning results with taxonomic classification, GC content, coverage, and tetranucleotide signatures [7]. The refined taxonomic information of assembled contigs were revealed precisely by Kaiju (ver. 1.7.3) against the NCBI nucleotide database (nr/nt). Gene models were predicted by Prodigal (ver. 2.6.3) [8] and genes were annotated using HMMer, ver. 3.1b2 [9] with Pfams, ver. 33.1 [10] and TIGRFAMs ver. 15.0 [11]. To further improve binning quality and to generate high-confidence reconstructed genomes, we manually curated bins using Anvi’o (ver. 6.2) to visualize and refine metagenomic bins [12]. Strain-level relationships were determined using StrainPhlAn (v 3.0) [13]. Genome completeness and redundancy of curated MAGs was checked with Anvi’o and all had redundancy below 10%. The manually curated MAG quality information for each of the 8 consortia is presented in Supplementary Table 1.

CAZyme gene annotation was performed using HMMer with the dbCAN2 database, ver. V8 [14] as a reference. CAZyme gene sequences of GH10 or GH43 from all MAGs were pooled, and then identical sequences were merged. 39 unique sequences were identified in GH10 gene pools, whereas 823 unique sequences were in GH43 pools. Unique sequences in GH10 or GH43 pools were aligned separately using Muscle v5 with super5 algorithm [15]. Maximum-likelihood trees were built using FastTree (2.1.11) with the JTT+CAT model [16]. Trees were visualized with annotations by iToL (v5) [17].

**Measurement of SCFAs**

Culture (1 mL) from each sample was centrifuged (18,000 × *g*, 10 min) to remove bacterial cells and the 400 μL of supernatant was transferred into a glass autosampler vial (Dwk Life Sciences LLC, Rockwood, TN) with 100 μL internal standards. Internal standards were prepared with 4-methylvaleric acid, phosphoric acid, and copper sulfate pentahydrate. SCFAs were quantified on an Agilent 7890A gas chromatograph (GC-FID 7890A, Santa Clara, CA) with a fused silica capillary column (Nukol, Supelco 40369-03A, 30 m × 0.25 mm, id 0.25 μm, Palo Alto, CA, USA). Equipment conditions: injector temperature at 230 °C; injection volume of 1 uL with split inlet, initial oven temperature at 100 °C; temperature ramp of 8 °C per min to 200 °C with a hold for 3 min at final stage. Helium was used as the carrier gas at 0.75 ml/min.

**Measurement of other polar metabolites**

Water extractable metabolites from day 7 cultures, including lactic acid, citric acid, formic acid, pyruvic acid, malic acid, succinic acid and fumaric acid were measured on a HPLC system (Waters Corporation, Milford, MA) with an organic acid column (BioRad Aminex HPX-87) and a refractive detector (Model 2414, Waters Corporation, Milford, MA).

**Bligh-Dyer extraction and trimethylsilyl derivatization**

Final cultures (day 7) of each SAX fermentation were subjected to a Bligh-Dyer extraction to extract polar metabolites. Briefly, 100 μL of culture supernatant was mixed with 300 μL of MeOH/CHCl_3_/H_2_O (1:1:1) solution. After 30 min incubation at 4 °C, samples were centrifuged (13,000 × *g*, 5 min). The aqueous fractions were collected and then dried with a centrifuge evaporator. The Bligh-Dyer extracted samples were treated with a trimethylsilyl (TMS) derivatization protocol. Methoxyamine hydrochloride (#226904, Sigma-Aldrich Co.) was dissolved in pure pyridine (#270407, Sigma-Aldrich Co.) to a final concentration of 20 mg/mL. Then, 40 μL of the fresh prepared methoxyamination reagent was added to the dried sample pellets. Samples were incubated at 37 °C for 2 h with continuous shaking (150 rpm). After incubation, 70 μL silylation reagent (N-Methyl-N-trimethylsilyl-trifluoroacetamide, #69479, Sigma-Aldrich Co.) was added with additional incubation for 30 min. The derivatized samples were then measured for non-targeted metabolomics by a Thermo Trace 1310 GC (Thermo Fisher Scientific Inc., Waltham, MA) coupled with Thermo TSQ 8000 triple quadrupole mass spectrometer (Thermo Fisher Scientific Inc., Waltham, MA) at the Purdue Metabolite Profiling Facility.

**Statistical analysis**

Statistical analyses were conducted on triplicated data. Gas, pH, α-diversity, OTU abundance and SCFAs data are presented as means and standard deviation. All statistical differences were calculated by analysis of variance (ANOVA). Tukey’s test with α level of 0.05 was adopted to differentiate significant mean variances. Statistical tests were conducted with R 4.1.0 (R Foundation for Statistical Computing, Vienna, Austria).

**References for supplementary information**

1.	Thakkar RD, Tuncil YE, Hamaker BR, Lindemann SR. Maize Bran Particle Size Governs the Community Composition and Metabolic Output of Human Gut Microbiota in in vitro Fermentations. *Front Microbiol* 2020; **11**.

2.	Quast C, Pruesse E, Yilmaz P, Gerken J, Schweer T, Yarza P, et al. The SILVA ribosomal RNA gene database project: improved data processing and web-based tools. *Nucleic Acids Res* 2013; **41**: D590–D596.

3.	Cole JR, Wang Q, Fish JA, Chai B, McGarrell DM, Sun Y, et al. Ribosomal Database Project: data and tools for high throughput rRNA analysis. *Nucleic Acids Res* 2014; **42**: D633–D642.

4.	Tuncil YE, Thakkar RD, Marcia ADR, Hamaker BR, Lindemann SR. Divergent short-chain fatty acid production and succession of colonic microbiota arise in fermentation of variously-sized wheat bran fractions. *Sci Rep* 2018; **8**: 16655.

5.	Gurevich A, Saveliev V, Vyahhi N, Tesler G. QUAST: quality assessment tool for genome assemblies. *Bioinforma Oxf Engl* 2013; **29**: 1072–1075.

6.	Wu Y-W, Simmons BA, Singer SW. MaxBin 2.0: an automated binning algorithm to recover genomes from multiple metagenomic datasets. *Bioinformatics* 2016; **32**: 605–607.

7.	Parks DH, Rinke C, Chuvochina M, Chaumeil P-A, Woodcroft BJ, Evans PN, et al. Recovery of nearly 8,000 metagenome-assembled genomes substantially expands the tree of life. *Nat Microbiol* 2017; **2**: 1533–1542.

8.	Hyatt D, Chen G-L, LoCascio PF, Land ML, Larimer FW, Hauser LJ. Prodigal: prokaryotic gene recognition and translation initiation site identification. *BMC Bioinformatics* 2010; **11**: 119.

9.	Johnson LS, Eddy SR, Portugaly E. Hidden Markov model speed heuristic and iterative HMM search procedure. *BMC Bioinformatics* 2010; **11**: 431.

10.	Finn RD, Bateman A, Clements J, Coggill P, Eberhardt RY, Eddy SR, et al. Pfam: the protein families database. *Nucleic Acids Res* 2014; **42**: D222–D230.

11.	Haft DH, Selengut JD, White O. The TIGRFAMs database of protein families. *Nucleic Acids Res* 2003; **31**: 371–373.

12.	Eren AM, Esen ÖC, Quince C, Vineis JH, Morrison HG, Sogin ML, et al. Anvi’o: an advanced analysis and visualization platform for ‘omics data. *PeerJ* 2015; **3**: e1319.

13.	Beghini F, McIver LJ, Blanco-Míguez A, Dubois L, Asnicar F, Maharjan S, et al. Integrating taxonomic, functional, and strain-level profiling of diverse microbial communities with bioBakery 3. *eLife* 2021; **10**: e65088.

14.	Zhang H, Yohe T, Huang L, Entwistle S, Wu P, Yang Z, et al. dbCAN2: a meta server for automated carbohydrate-active enzyme annotation. *Nucleic Acids Res* 2018; **46**: W95–W101.

15.	Edgar RC. MUSCLE: multiple sequence alignment with high accuracy and high throughput. *Nucleic Acids Res* 2004; **32**: 1792–1797.

16.	Price MN, Dehal PS, Arkin AP. FastTree: Computing Large Minimum Evolution Trees with Profiles instead of a Distance Matrix. *Mol Biol Evol* 2009; **26**: 1641–1650.

17.	Letunic I, Bork P. Interactive Tree Of Life (iTOL) v5: an online tool for phylogenetic tree display and annotation. *Nucleic Acids Res* 2021; **49**: W293–W296.
