## Supplementary material for "SUBTLE DIFFERENCES IN FINE POLYSACCHARIDE STRUCTURES GOVERN SELECTION AND SUCCESSION OF HUMAN GUT MICROBIOTA": Table S1

Table S1. Reconstructed genome bins from shotgun sequencing of day7 consortia ^A^

| RSAX donor3 | Taxonomy identification | Bin abundance (%) | Completion (%) | Redundancy (%) |
| --- | --- | --- | --- | --- |
| Bin01 | *Agathobacter rectale* | 14.74 | 71.83 | 8.45 |
| Bin02 | *Lachnospiraceae GCAsp900066135* | 11.92 | 98.59 | 7.04 |
| Bin03 | *Bifidobacterium longum* | 8.78 | 97.18 | 0.00 |
| Bin04 | *Eisenbergiella tayi* | 7.13 | 100.00 | 2.82 |
| Bin05 | *Blautia producta* | 6.02 | 100.00 | 0.00 |
| Bin06 | *Bacteroides ovatus* | 5.99 | 88.73 | 4.23 |
| Bin07 | *Escherichia coli* | 4.68 | 100.00 | 0.00 |
| Bin08 | *Fusicatenibacter saccharivorans* | 4.23 | 98.59 | 2.82 |
| Bin09 | *Bacteroides xylanisolvens* | 4.23 | 0.00 | 0.00 |
| Bin10 | *Blautia wexlerae* | 3.19 | 97.18 | 4.23 |
| Bin11 | *Bacteroides fragilis* | 2.02 | 73.24 | 4.23 |
| Bin12 | *Bacteroides dorei* | 1.42 | 81.69 | 4.23 |
| Bin13 | *Murimonas intestini* | 1.33 | 92.96 | 7.04 |
| Bin14 | *Bifidobacterium adolescentis* | 1.30 | 69.01 | 2.82 |
| Bin15 | *Anaerostipes hadrus* | 0.93 | 98.59 | 4.23 |
| Bin16 | *Lachnospiraceae CAGsp900066615* | 0.72 | 100.00 | 4.23 |
| Bin17 | *Monoglobalas UGAsp900066215* | 0.70 | 100.00 | 0.00 |
| Bin18 | *Agathobaculum butyriciproducens* | 0.68 | 87.32 | 5.63 |
| Bin19 | *Lachnospiraceae TF11sp436755* | 0.62 | 100.00 | 5.63 |
| Bin20 | *Roseburia intestinalis* | 0.57 | 94.37 | 7.04 |
| Bin21 | *Eisenbergiella massiliensis* | 0.45 | 53.52 | 4.23 |
| Bin22 | *Acutalibacteraceae UBAsp3531055* | 0.39 | 98.59 | 4.23 |
| Bin23 | *Eubacterium sp000434315* | 0.38 | 97.18 | 11.27 |
| Bin24 | *Bacteroides stercoris* | 0.37 | 0.00 | 0.00 |
| Bin25 | *Erysipelatoclostridium ramosum* | 0.36 | 97.18 | 0.00 |
| Bin26 | *Phascolarctobacterium faecium* | 0.35 | 100.00 | 1.41 |
| Bin27 | *Lachnospiraceae CGA303sp437755* | 0.34 | 100.00 | 1.41 |
| Bin28 | *Desulfovibrio piger* | 0.33 | 95.77 | 1.41 |
| Bin29 | *Ruminococcus bicirulans* | 0.31 | 100.00 | 5.63 |
| Bin30 | *Lachnospira sp436475* | 0.27 | 77.46 | 8.45 |
| Bin31 | *Hungatella hathewayi* | 0.26 | 88.73 | 4.23 |
| Bin32 | *Faecalibacterium prausnitziiD* | 0.26 | 50.70 | 8.45 |
| Bin33 | *Agathobaculum butyriciproducens2* | 0.22 | 54.93 | 1.41 |
| Bin34 | *Blautia obeum* | 0.22 | 0.00 | 0.00 |
| Bin35 | *Anaerotignum lactatifermentans* | 0.21 | 91.55 | 4.23 |
| Bin36 | *Eubacterium hallii* | 0.20 | 50.70 | 0.00 |
| Bin37 | *Clostridium bolteae* | 0.18 | 56.34 | 4.23 |
| Bin38 | *Anaeromassilibacillus sp1305115* | 0.17 | 67.61 | 1.41 |
| Bin39 | *Lachnospiraceae UBAsp2160135* | 0.16 | 47.89 | 1.41 |
| Bin40 | *Marseille sp900232885* | 0.14 | 47.89 | 0.00 |
| Bin41 | *Oscillibacter sp001916835* | 0.13 | 0.00 | 0.00 |
| Bin42 | *Dakarella massiliensis* | 0.12 | 52.11 | 0.00 |
| Unbinned | *Unbinned contigs* | 15.56 | N/A | N/A |
| RSAX donor1 | Taxonomy identification | Bin abundance (%) | Completion (%) | Redundancy (%) |
| Bin01 | *Agathobacter rectale* | 22.56 | 98.59 | 7.04 |
| Bin02 | *Lachnospiraceae GCAsp900066135* | 13.10 | 88.73 | 0.00 |
| Bin03 | *Blautia producta* | 8.26 | 100.00 | 1.41 |
| Bin04 | *Eisenbergiella tayi* | 5.75 | 100.00 | 2.82 |
| Bin05 | *Bacteroides ovatus* | 4.31 | 50.70 | 8.45 |
| Bin06 | *Bacteroides xylanisolvens* | 3.61 | 0.00 | 0.00 |
| Bin07 | *Blautia wexlerae* | 3.55 | 97.18 | 4.23 |
| Bin08 | *Escherichia coli* | 3.52 | 100.00 | 0.00 |
| Bin09 | *Fusicatenibacter saccharivorans* | 2.63 | 98.59 | 1.41 |
| Bin10 | *Bacteroides stercoris* | 1.70 | 63.38 | 4.23 |
| Bin11 | *Ruminococcus bicirculans* | 1.69 | 100.00 | 4.23 |
| Bin12 | *Bacteroides dorei* | 1.61 | 94.37 | 1.41 |
| Bin13 | *Bifidobacterium longum* | 1.24 | 98.59 | 0.00 |
| Bin14 | *Bifidobacterium pseudocatenulatum* | 1.24 | 98.59 | 1.41 |
| Bin15 | *Phascolarctobacterium succinatutens* | 0.75 | 100.00 | 4.23 |
| Bin16 | *Eubacterium sp000432355* | 0.73 | 45.07 | 2.82 |
| Bin17 | *Monoglobales CAG41sp900066215* | 0.72 | 100.00 | 1.41 |
| Bin18 | *Lachnospiraceae CAGsp900066615* | 0.69 | 100.00 | 8.45 |
| Bin19 | *Blautia obeum* | 0.64 | 73.24 | 8.45 |
| Bin20 | *Eisenbergiella massiliensis* | 0.52 | 57.75 | 2.82 |
| Bin21 | *Roseburia intestinalis* | 0.52 | 84.51 | 7.04 |
| Bin22 | *Lachnospiraceae CAGsp437755* | 0.50 | 100.00 | 1.41 |
| Bin23 | *Anaerostipes hadrus* | 0.47 | 98.59 | 4.23 |
| Bin24 | *Agathobaculum butyriciproducens* | 0.40 | 92.96 | 9.86 |
| Bin25 | *Hungatella hathewayi* | 0.36 | 87.32 | 8.45 |
| Bin26 | *Erysipelatoclostridium ramosum* | 0.34 | 97.18 | 0.00 |
| Bin27 | *Lachnospira sp000436475* | 0.34 | 100.00 | 8.45 |
| Bin28 | *Agathobaculum sp900291975* | 0.33 | 64.79 | 0.00 |
| Bin29 | *Oscillibacter sp001916835* | 0.25 | 94.37 | 7.04 |
| Bin30 | *Bacteroides uniformis* | 0.24 | 50.70 | 2.82 |
| Bin31 | *Eubacterium hallii* | 0.22 | 95.77 | 5.63 |
| Bin32 | *Eubacterium sp000435815* | 0.20 | 100.00 | 0.00 |
| Bin33 | *Lachnospiraceae UBAsp2160135* | 0.18 | 94.37 | 5.63 |
| Bin34 | *Oxalobacter formigenes* | 0.17 | 95.77 | 0.00 |
| Bin35 | *Marseille sp900232885* | 0.17 | 35.21 | 1.41 |
| Bin36 | *Parabacteroides johnsonii* | 0.16 | 90.14 | 7.04 |
| Bin37 | *Anaerotignaceae An114sp2161055* | 0.15 | 91.55 | 1.41 |
| Bin38 | *Anaerofustis stercorihominis* | 0.15 | 94.37 | 1.41 |
| Bin39 | *Dakarella massiliensis* | 0.13 | 88.73 | 2.82 |
| Bin40 | *Clostridium lavalense* | 0.12 | 57.75 | 1.41 |
| Bin41 | *Acutalibacteraceae UBAsp3531055* | 0.11 | 61.97 | 0.00 |
| Bin42 | *Lactonifactor longoviformis* | 0.10 | 0.00 | 0.00 |
| Unbinned | *Unbinned contigs* | 12.98 | N/A | N/A |
| RSAX donor2 | Taxonomy identification | Bin abundance (%) | Completion (%) | Redundancy (%) |
| Bin01 | *Bacteroides xylanisolvens* | 18.10 | 50.70 | 4.23 |
| Bin02 | *Bacteroides ovatus* | 17.84 | 33.80 | 1.41 |
| Bin03 | *Bifidobacterium longum* | 15.49 | 100.00 | 1.41 |
| Bin04 | *Esenbergiella tayi* | 11.85 | 100.00 | 2.82 |
| Bin05 | *Escherichia coli* | 6.46 | 100.00 | 0.00 |
| Bin06 | *Bifidobacterium adolescentis* | 2.97 | 74.65 | 2.82 |
| Bin07 | *Blautia sp001304935* | 2.23 | 0.00 | 0.00 |
| Bin08 | *Blautia sp900066165* | 1.77 | 97.18 | 5.63 |
| Bin09 | *Anaerostipes sp000508985* | 1.53 | 100.00 | 5.63 |
| Bin10 | *Agathobaculum butyriciproducens* | 1.06 | 100.00 | 4.23 |
| Bin11 | *Blautia wexlerae* | 1.04 | 98.59 | 4.23 |
| Bin12 | *Clostridium bolteae* | 0.86 | 80.28 | 1.41 |
| Bin13 | *Bacteroides uniformis* | 0.65 | 78.87 | 5.63 |
| Bin14 | *Clostridium symbiosum* | 0.65 | 88.73 | 0.00 |
| Bin15 | *Blautia hominis* | 0.65 | 98.59 | 8.45 |
| Bin16 | *Parabacteroides goldsteinii* | 0.59 | 69.01 | 2.82 |
| Bin17 | *Parabacteroides timonensis* | 0.57 | 33.80 | 1.41 |
| Bin18 | *Monoglobale CAG41sp900066215* | 0.54 | 100.00 | 0.00 |
| Bin19 | *Parabacteroides merdae* | 0.42 | 77.46 | 2.82 |
| Bin20 | *Blautia producta* | 0.42 | 0.00 | 0.00 |
| Bin21 | *Clostridium sp000155435* | 0.39 | 57.75 | 0.00 |
| Bin22 | *Lachnospiraceae KLEsp66985* | 0.39 | 98.59 | 7.04 |
| Bin23 | *Clostridium citroniae* | 0.38 | 60.56 | 4.23 |
| Bin24 | *Anaerotignaceae An114sp3543235* | 0.33 | 92.96 | 4.23 |
| Bin25 | *Erysipelatoclostridium sp003024675* | 0.31 | 97.18 | 11.27 |
| Bin26 | *Bacteroides stercoris* | 0.29 | 40.85 | 1.41 |
| Bin27 | *Klebsiella variicola* | 0.27 | 0.00 | 0.00 |
| Bin28 | *Hungatella hathewayi* | 0.24 | 0.00 | 0.00 |
| Bin29 | *Erysipelatoclostridium ramosum* | 0.21 | 42.25 | 0.00 |
| Unbinned | *Unbinned contigs* | 11.51 | N/A | N/A |
| RSAX pooled | Taxonomy identification | Bin abundance (%) | Completion (%) | Redundancy (%) |
| Bin01 | *Agathobacter rectale* | 19.42 | 98.59 | 7.04 |
| Bin02 | *Murimonas intestini* | 13.05 | 100.00 | 1.41 |
| Bin03 | *Bifidobacterium longum* | 12.61 | 100.00 | 0.00 |
| Bin04 | *Blautia wexlerae* | 7.67 | 98.59 | 4.23 |
| Bin05 | *Eisenbergiella tayi* | 5.88 | 92.96 | 5.63 |
| Bin06 | *Fusicatenibacter saccharivorans* | 5.28 | 97.18 | 2.82 |
| Bin07 | *Escherichia coli* | 3.79 | 100.00 | 0.00 |
| Bin08 | *Bacteroides fragilis* | 3.32 | 94.37 | 2.82 |
| Bin09 | *Bacteroides ovatus* | 2.52 | 98.59 | 5.63 |
| Bin10 | *Desulfovibrio piger* | 2.46 | 98.59 | 1.41 |
| Bin11 | *Bacteroides uniformis* | 1.65 | 97.18 | 2.82 |
| Bin12 | *Bacteroides vulgatus* | 1.43 | 98.59 | 2.82 |
| Bin13 | *Clostridium bolteae* | 1.15 | 84.51 | 8.45 |
| Bin14 | *Eisenbergiella massiliensis* | 1.14 | 63.38 | 4.23 |
| Bin15 | *Lachnospiraceae TFsp436755* | 1.00 | 100.00 | 4.23 |
| Bin16 | *Parabacteroides johnsonii* | 0.97 | 100.00 | 5.63 |
| Bin17 | *Monoglobales CAGsp900066215* | 0.52 | 100.00 | 1.41 |
| Bin18 | *Eubacterium sp000435815* | 0.43 | 100.00 | 5.63 |
| Bin19 | *Coprococcus comes* | 0.37 | 100.00 | 5.63 |
| Bin20 | *Lachnospiraceae CAGsp900066615* | 0.32 | 95.77 | 1.41 |
| Bin21 | *Erysipelatoclostridium ramosum* | 0.32 | 97.18 | 0.00 |
| Bin22 | *Lachnospira sp900316325* | 0.28 | 90.14 | 5.63 |
| Bin23 | *Clostridium symbiosum* | 0.26 | 90.14 | 0.00 |
| Bin24 | *Bacteroides thetaiotaomicron* | 0.25 | 40.85 | 2.82 |
| Bin25 | *Phascolarctobacterium faecium* | 0.25 | 100.00 | 1.41 |
| Bin26 | *Lachnospiracea KLEsp900066985* | 0.20 | 95.77 | 1.41 |
| Bin27 | *Bifidobacterium pseudocatenulatum* | 0.20 | 64.79 | 0.00 |
| Bin28 | *Faecalibacterium prausnitziiG* | 0.19 | 61.97 | 4.23 |
| Bin29 | *Bacteroides timonensis* | 0.19 | 47.89 | 5.63 |
| Bin30 | *Clostridium sp000155435* | 0.17 | 49.30 | 1.41 |
| Bin31 | *Acutalibacteraceae UBAsp3531055* | 0.17 | 97.18 | 5.63 |
| Bin32 | *Faecalibacterium prausnitziiJ* | 0.16 | 63.38 | 9.86 |
| Bin33 | *Massilioclostridium methylpentosum* | 0.15 | 73.24 | 0.00 |
| Bin34 | *Lachnospira sp000437735* | 0.14 | 85.92 | 5.63 |
| Bin35 | *Anaerotignaceae An114sp3531055* | 0.13 | 91.55 | 0.00 |
| Bin36 | *Agathobaculum butyriciproducens* | 0.12 | 77.46 | 0.00 |
| Bin37 | *Bacteroides eggerthii* | 0.12 | 19.72 | 1.41 |
| Bin38 | *Oxalobacter formigenes* | 0.11 | 70.42 | 2.82 |
| Bin39 | *Dorea sp000509125* | 0.11 | 53.52 | 8.45 |
| Bin40 | *Lachnospiracea UBAsp2160135* | 0.11 | 66.20 | 1.41 |
| Bin41 | *Ruminiclostridium siraeum* | 0.10 | 61.97 | 4.23 |
| Bin42 | *Hungatella effluvii* | 0.09 | 23.94 | 0.00 |
| Unbinned | *Unbinned contigs* | 11.21 | N/A | N/A |
| WSAX donor1 | Taxonomy identification | Bin abundance (%) | Completion (%) | Redundancy (%) |
| Bin01 | *Agathobacter rectale* | 26.91 | 98.59 | 7.04 |
| Bin02 | *Blautia producta* | 13.35 | 100.00 | 1.41 |
| Bin03 | *Blautia wexlerae* | 10.86 | 100.00 | 4.23 |
| Bin04 | *Bifidobacterium longum* | 7.33 | 98.59 | 0.00 |
| Bin05 | *Bifidobacterium pseudocatenulatum* | 6.71 | 98.59 | 0.00 |
| Bin06 | *Escherichia coli* | 5.61 | 100.00 | 0.00 |
| Bin07 | *Faecalibacterium prausnitziiA* | 2.46 | 81.69 | 7.04 |
| Bin08 | *Faecalibacterium prausnitziiE* | 2.26 | 92.96 | 5.63 |
| Bin09 | *Bacteroides dorei* | 1.84 | 100.00 | 2.82 |
| Bin10 | *Coprococcus comes* | 1.13 | 100.00 | 7.04 |
| Bin11 | *Bacteroides xylanisolvens* | 0.74 | 0.00 | 0.00 |
| Bin12 | *Bacteroides ovatus* | 0.54 | 0.00 | 0.00 |
| Bin13 | *Erysipelatoclostridium ramosum* | 0.50 | 97.18 | 0.00 |
| Bin14 | *Phascolarctobacterium succinatutens* | 0.48 | 100.00 | 1.41 |
| Bin15 | *Bacteroides stercoris* | 0.35 | 67.61 | 1.41 |
| Bin16 | *Gemmiger formicilis* | 0.31 | 78.87 | 1.41 |
| Bin17 | *Agathobaculum butyriciproducens* | 0.31 | 100.00 | 4.23 |
| Bin18 | *Bacteroides thetaiotaomicron* | 0.28 | 56.34 | 2.82 |
| Bin19 | *Gemmiger formicilis* | 0.14 | 0.00 | 0.00 |
| Bin20 | *Hungatella effluvii* | 0.13 | 53.52 | 2.82 |
| Unbinned | *Unbinned contigs* | 17.73 | N/A | N/A |
| WSAX donor2 | Taxonomy identification | Bin abundance (%) | Completion (%) | Redundancy (%) |
| Bin01 | *Agathobacter rectale* | 29.59 | 98.59 | 7.04 |
| Bin02 | *Bifidobacterium longum* | 25.06 | 100.00 | 0.00 |
| Bin03 | *Bifidobacterium adolescentis* | 8.08 | 92.96 | 0.00 |
| Bin04 | *Bifidobacterium dentium* | 6.49 | 98.59 | 2.82 |
| Bin05 | *Escherichia coli* | 5.97 | 100.00 | 0.00 |
| Bin06 | *Blautia hominis* | 2.90 | 100.00 | 0.00 |
| Bin07 | *Bacteroides xylanisolvens* | 2.44 | 84.51 | 2.82 |
| Bin08 | *Bacteroides ovatus* | 2.09 | 39.44 | 2.82 |
| Bin09 | *Hungatella hathewayi* | 1.93 | 95.77 | 7.04 |
| Bin10 | *Erysipelatoclostridium ramosum* | 0.74 | 97.18 | 0.00 |
| Bin11 | *Eisenbergiella tayi* | 0.73 | 100.00 | 2.82 |
| Bin12 | *Faecalibacterium prausnitziiG* | 0.37 | 95.77 | 5.63 |
| Bin13 | *Bacteroides uniformis* | 0.34 | 84.51 | 4.23 |
| Bin14 | *Enterococcus Dsp002850555* | 0.32 | 100.00 | 4.23 |
| Bin15 | *Bacteroides thetaiotaomicron* | 0.30 | 0.00 | 0.00 |
| Bin16 | *Blautia wexlerae* | 0.29 | 100.00 | 4.23 |
| Bin17 | *Phascolarctobacterium faecium* | 0.29 | 100.00 | 1.41 |
| Bin18 | *Faecalicatena gnavus* | 0.27 | 100.00 | 0.00 |
| Bin19 | *Bacteroides dorei* | 0.19 | 43.66 | 2.82 |
| Bin20 | *Bacteroides vulgatus* | 0.18 | 0.00 | 0.00 |
| Bin21 | *Lachnospiraceae KLEsp900066985* | 0.17 | 100.00 | 1.41 |
| Bin22 | *Anaerostipes sp508985* | 0.16 | 97.18 | 4.23 |
| Bin23 | *Veillonella parvula* | 0.07 | 94.37 | 1.41 |
| Unbinned | *Unbinned contigs* | 11.01 | N/A | N/A |
| WSAX donor3 | Taxonomy identification | Bin abundance (%) | Completion (%) | Redundancy (%) |
| Bin01 | *Bifidobacterium longum* | 21.84 | 100.00 | 0.00 |
| Bin02 | *Agathobacter rectale* | 18.36 | 98.59 | 5.63 |
| Bin03 | *Bacteroides fragilis* | 11.26 | 98.59 | 4.23 |
| Bin04 | *Hungatella effluvii* | 9.61 | 97.18 | 9.86 |
| Bin05 | *Eisenbergiella tayi* | 8.38 | 94.37 | 2.82 |
| Bin06 | *Escherichia coli* | 5.20 | 100.00 | 0.00 |
| Bin07 | *Bifidobacterium pseudocatenulatum* | 2.97 | 61.97 | 0.00 |
| Bin08 | *Lachnospiraceae TF01sp436755* | 2.19 | 100.00 | 4.23 |
| Bin09 | *Faecalibacterium prausnitziiD* | 1.20 | 60.56 | 14.08 |
| Bin10 | *Erysipelatoclostridium ramosum* | 0.82 | 97.18 | 0.00 |
| Bin11 | *Blautia sp900066165* | 0.74 | 97.18 | 1.41 |
| Bin12 | *Desulfovibrio piger* | 0.60 | 98.59 | 1.41 |
| Bin13 | *Coprococcus comes* | 0.24 | 98.59 | 5.63 |
| Bin14 | *Phascolarctobacterium faecium* | 0.21 | 100.00 | 1.41 |
| Bin15 | *Hungatella hathewayi* | 0.15 | 0.00 | 0.00 |
| Bin16 | *Lachnospiraceae GCAsp900066135* | 0.13 | 88.73 | 1.41 |
| Bin17 | *Blautia sp285855* | 0.10 | 50.70 | 4.23 |
| Bin18 | *Gemmiger formicilis* | 0.09 | 45.07 | 0.00 |
| Unbinned | *Unbinned contigs* | 15.91 | N/A | N/A |
| WSAX pooled | Taxonomy identification | Bin abundance (%) | Completion (%) | Redundancy (%) |
| Bin01 | *Agathobacter rectale* | 29.29 | 98.59 | 7.04 |
| Bin02 | *Bifidobacterium longum* | 25.88 | 100.00 | 0.00 |
| Bin03 | *Blautia producta* | 7.45 | 100.00 | 0.00 |
| Bin04 | *Bifidobacterium dentium* | 6.51 | 92.96 | 4.23 |
| Bin05 | *Escherichia coli* | 5.66 | 100.00 | 0.00 |
| Bin06 | *Bifidobacterium pseudocatenulatum* | 4.65 | 56.34 | 8.45 |
| Bin07 | *Faecalibacterium prausnitziiK* | 3.52 | 63.38 | 7.04 |
| Bin08 | *Bifidobacterium adolescentis* | 2.94 | 78.87 | 1.41 |
| Bin09 | *Coprococcus comes* | 1.00 | 100.00 | 7.04 |
| Bin10 | *Bacteroides fragilis* | 0.92 | 98.59 | 4.23 |
| Bin11 | *Hungatella hathewayi* | 0.90 | 95.77 | 7.04 |
| Bin12 | *Erysipelatoclostridium ramosum* | 0.80 | 97.18 | 0.00 |
| Bin13 | *Faecalibacterium prausnitziiE* | 0.32 | 54.93 | 7.04 |
| Bin14 | *Phascolarctobacterium faecium* | 0.25 | 100.00 | 1.41 |
| Bin15 | *Lachnospira eligens* | 0.25 | 100.00 | 5.63 |
| Bin16 | *Bacteroides ovatus* | 0.21 | 0.00 | 0.00 |
| Bin17 | *Bacteroides xylanisolvens* | 0.21 | 59.15 | 2.82 |
| Bin18 | *Bacteroides thetaiotaomicron* | 0.19 | 60.56 | 4.23 |
| Bin19 | *Bacteroides dorei* | 0.17 | 87.32 | 2.82 |
| Bin20 | *Gemmiger formicilis* | 0.13 | 97.18 | 0.00 |
| Bin21 | *Faecalibacterium prausnitziiD* | 0.13 | 0.00 | 0.00 |
| Bin22 | *Dakarella massiliensis* | 0.05 | 63.38 | 0.00 |
| Unbinned | *Unbinned contigs* | 8.59 | N/A | N/A |

^A^ Bin abundances were estimated by a proportion of bin coverages
