## Supplementary material for "SUBTLE DIFFERENCES IN FINE POLYSACCHARIDE STRUCTURES GOVERN SELECTION AND SUCCESSION OF HUMAN GUT MICROBIOTA": Supplimentary figures

Fig. S1

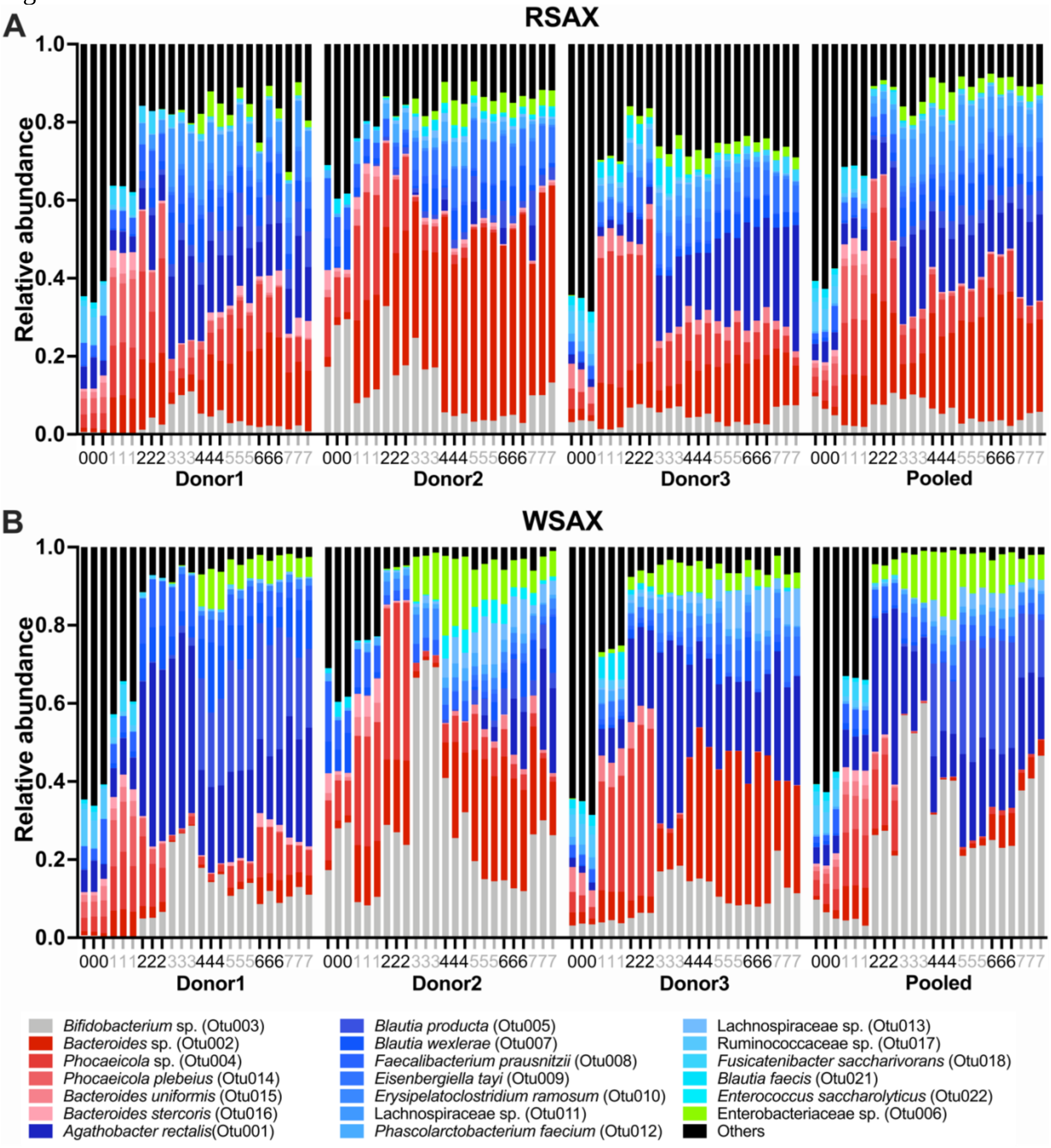


Fig. S1. Relative abundances of OTUs across all three replicates over 7 passages, as determined by 16S amplicon sequencing. X-axes represent passage numbers; number "0" denotes initial fecal inocula. Each shade represents a distinct OTU. They are colored by phylum (Pseudomonadota in green; Bacillota in blue; Bacteroidota in red and Actinomycetota in gray). Rare OTUs (< 0.2% relative abundance in total) are combined in "Others" (black).

Fig. S2

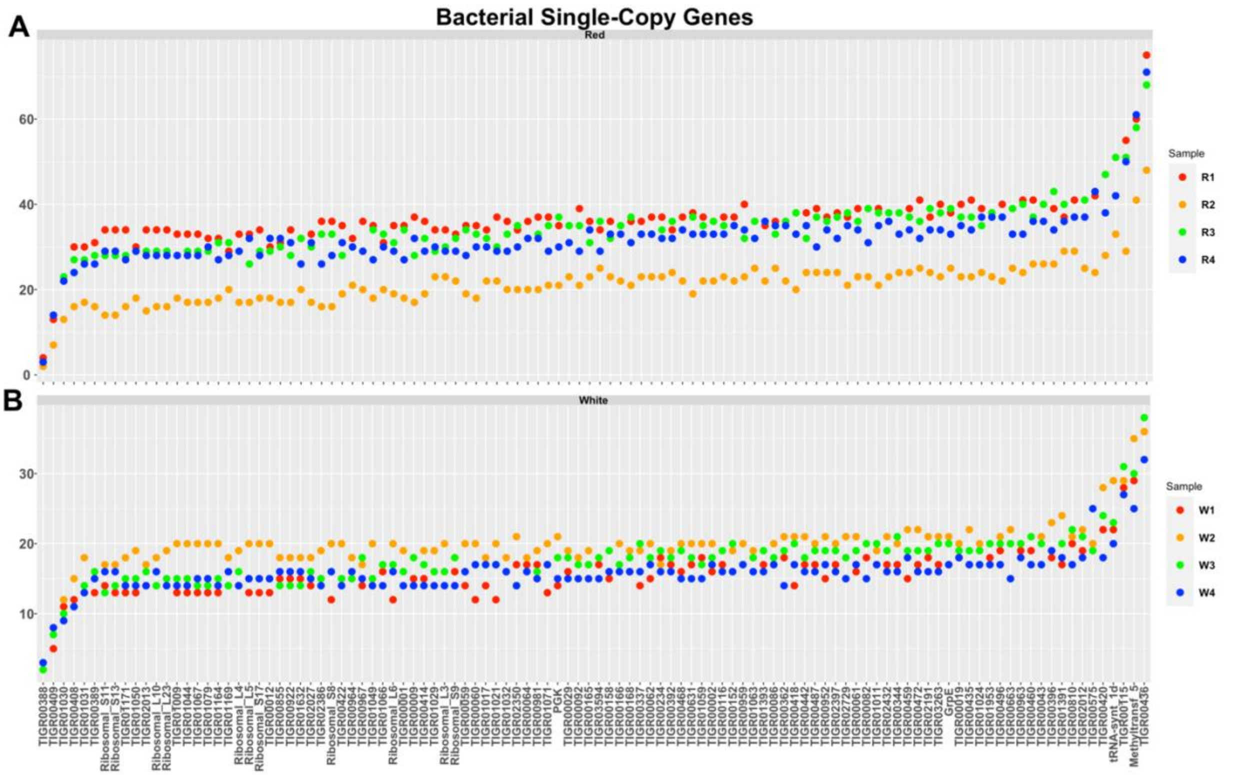


Fig. S2. Numbers of different bacterial single copy genes (BSCGs) in RSAX (A) and WSAX (B) consortia. Y-axis shows the counts of single-copy genes, and x-axis shows the TIGRfam family names of BSCGs. The numbers of BSCGs can indicate average diversity of observed bacterial organisms in each sample.

Fig. S3

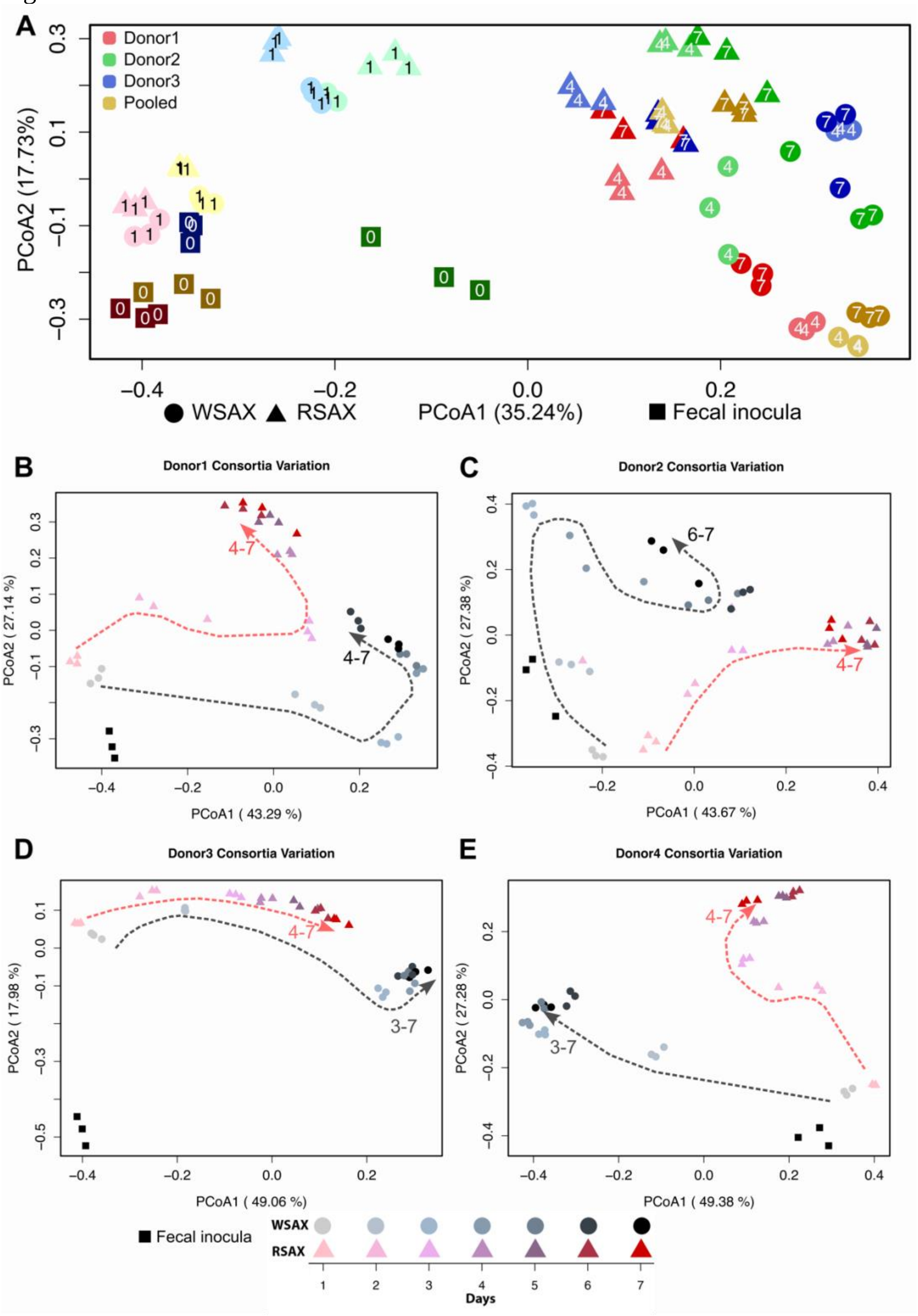


Fig. S3. Principal coordinate analysis (PCoA) plot with Bray-Curtis dissimilarity for microbial communities of fecal inocula, day 1, day 4 and day 7 (A). Numbers in each dot indicate different passage days. A clear separation between round dots (WSAX) and triangles (RSAX) was observed at later passages (day 4 and day 7). PCoA plot of Bray-Curtis dissimilarity across all sample lineages over 7 days (B-E). Different succession trajectories for Donor 1 (B), Donor 2 (C), Donor 3 (D) and pooled donor (E) inocula are illustrated with dashed lines. Numbers nearby arrows indicate passage days that clustered around final communities.

Fig. S4

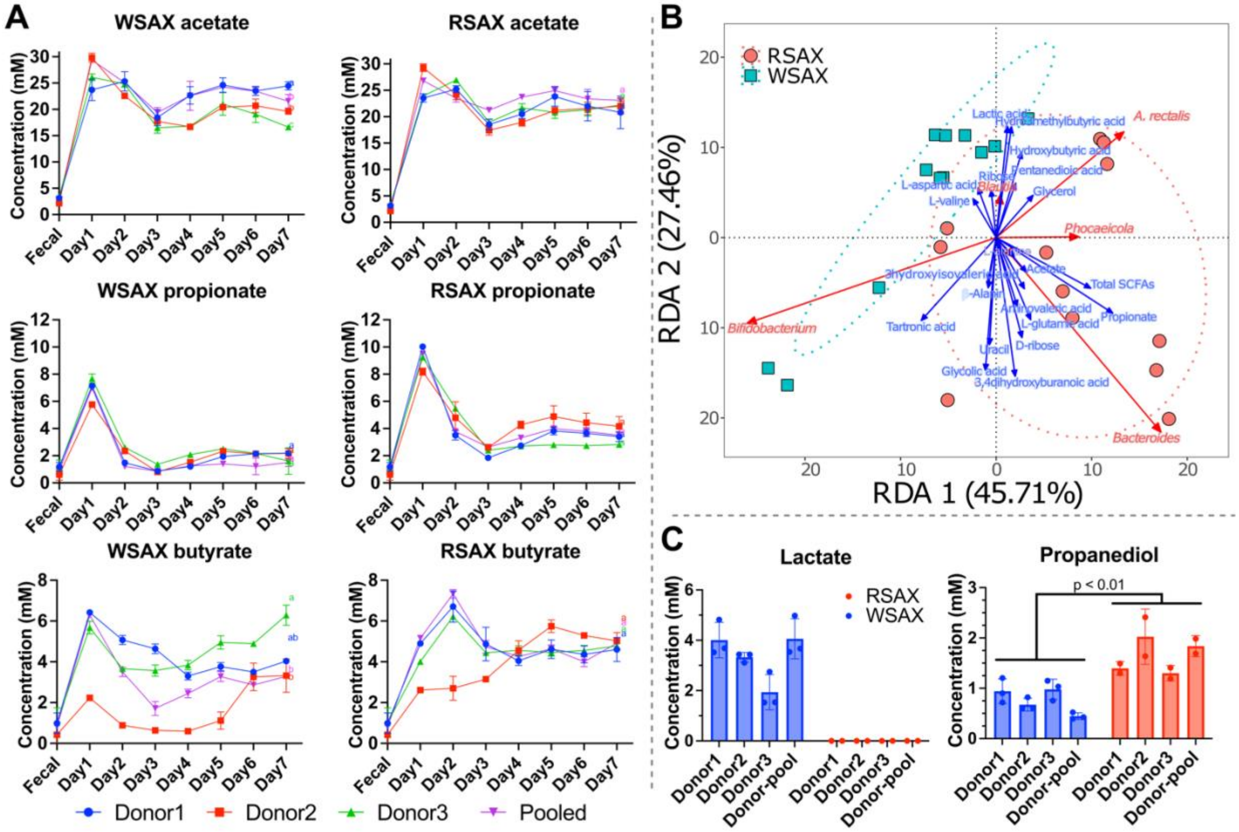


Fig. S4. Daily changes of SCFAs for WSAX and RSAX cultures (A). Different letters of each line indicate significant differences ($p < 0.05$) at day 7. Relevance of OTUs and metabolites in passage 7 communities, revealed by redundancy analysis (RDA) plot (B). HPLC quantification of general metabolites from day-7 cultures (C), with undetected succinate, pyruvate, maleic acid, formic acid, fumarate and methanol, whilst citric acid had non-differentiable values masked by buffer salts.

Fig. S5

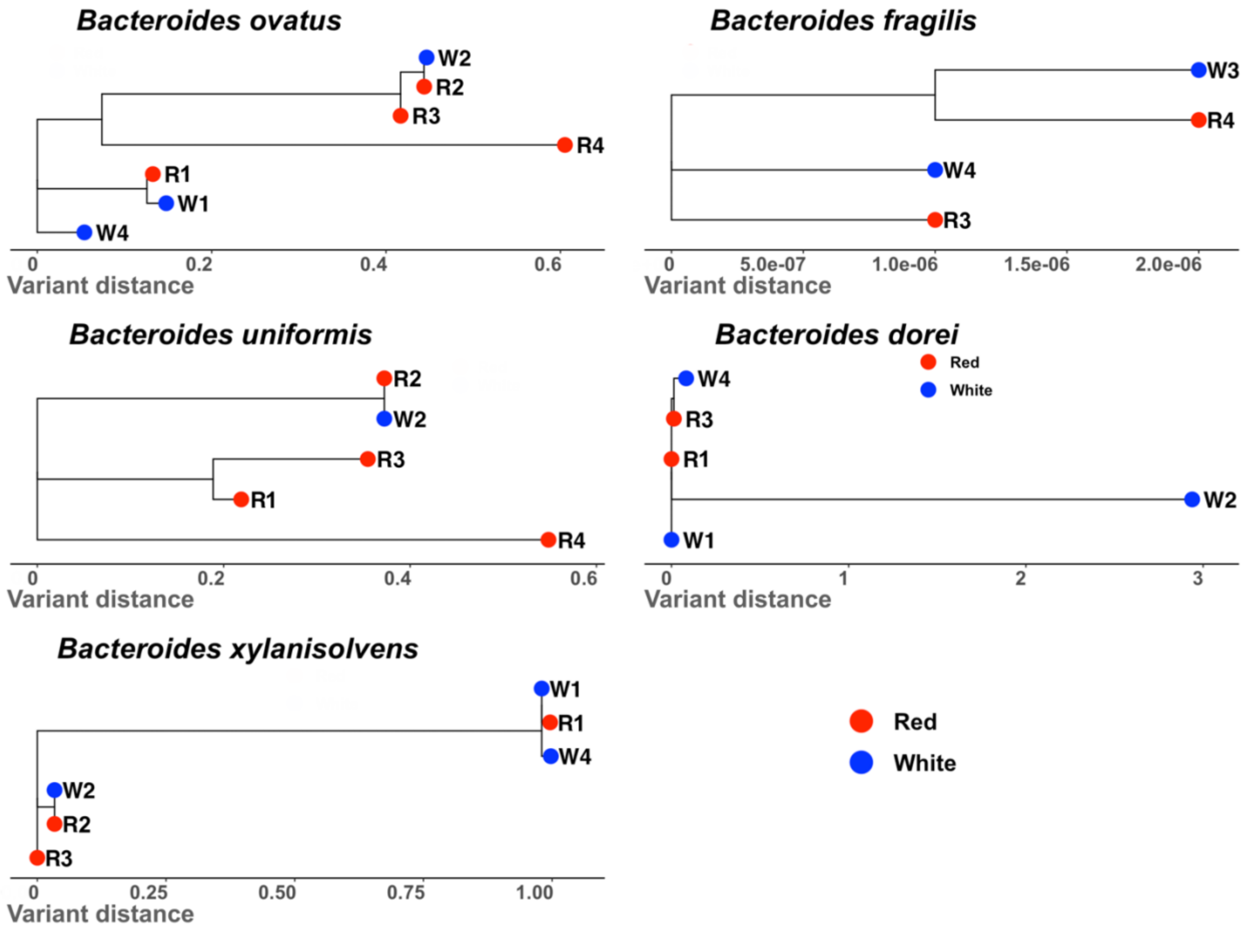


Fig. S5 Strain level differences from *Bacteroides* spp. in MAGs of final consortia. Different strain penetration among species showed evidence of donor-specificity and increased inter-strain competition. *B. ovatus* MAGs were recovered from each donors' consortium with different winning strains across RSAX and WSAX community. Similar patterns were observed for *B. uniformis*, *B. fragilis* and *B. xylanisolvens*. However, MAGs of *B. dorei* were nearly identical across different donors and only Donor 2 on WSAX had a divergent strain. Interestingly, MAGs not clustered with other donor-specific consortia were reconstructed from pooled consortia (*B. ovatus*, *B. uniformis*), suggesting that different *Bacteroides* spp. may be competitive under differing diversity conditions.

Fig. S6

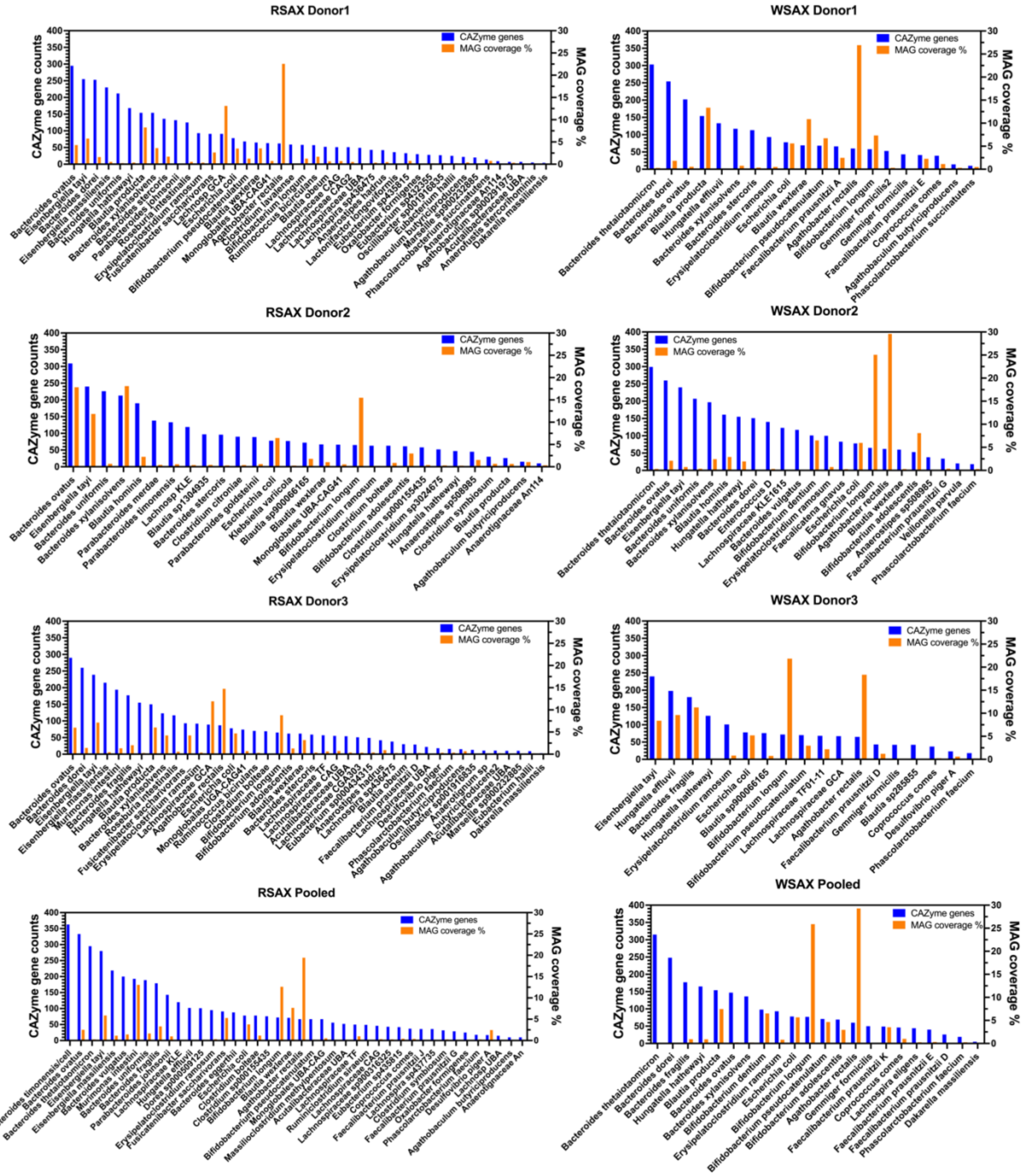


Fig. S6 Distribution of total numbers of CAZyme genes in each MAGs in comparison to MAG percent coverages. Non-unified distribution between CAZyme genes and MAG abundance suggests that high genetic potential of carbohydrate hydrolysis does not guarantee the high abundance in the competitive community.

Fig. S7

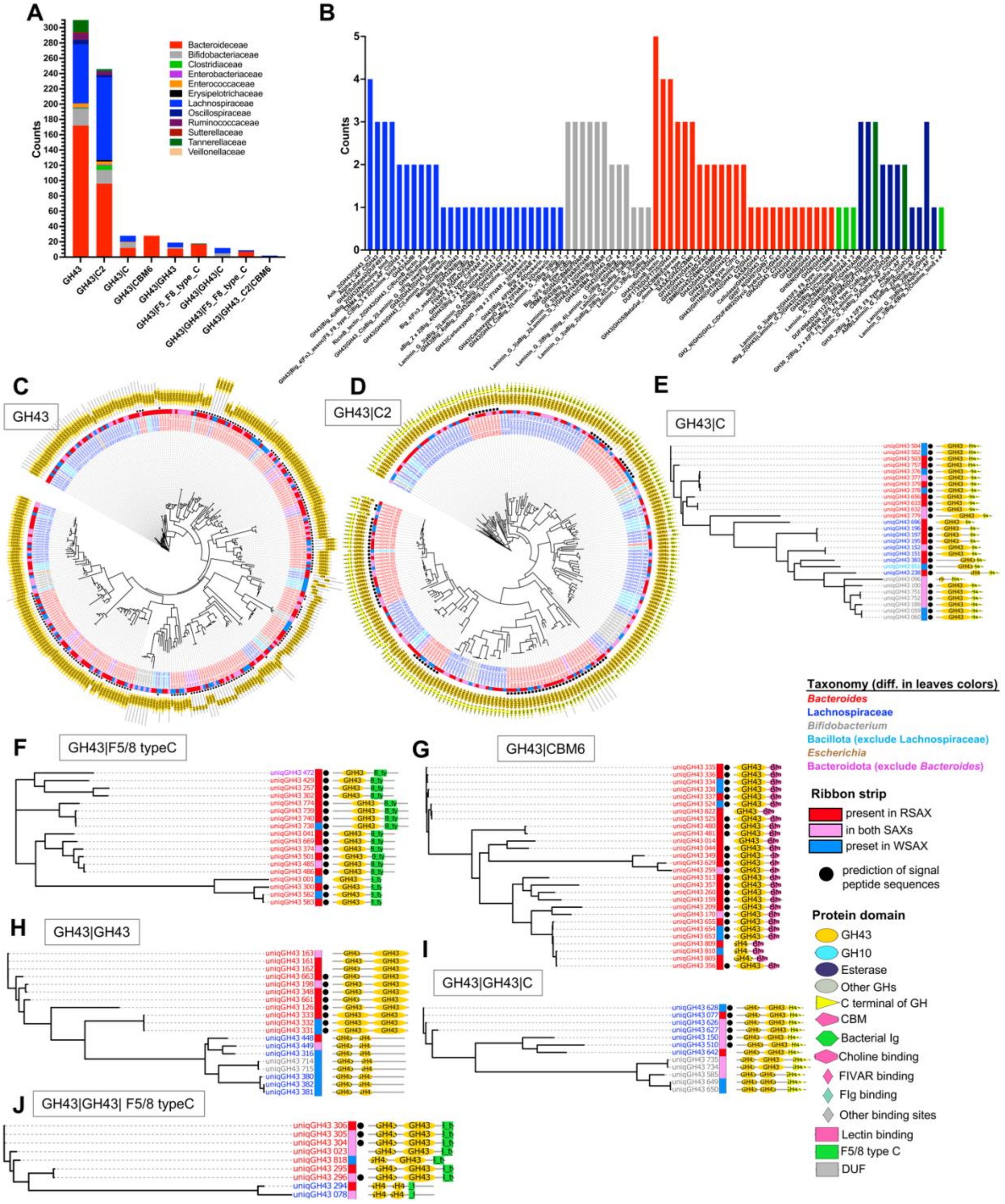


Fig. S7 Abundance distribution of AX-related GH43 at family-level taxonomy with simpler protein module structures (A) and relatively complicate domain structures (B). Phylogenetic trees of GH43 with simpler module structures (C-J). Labels of each leaf are colored at different taxonomy levels. Strips wrapped on leaves display substrate-related information. Black dots predict the presence of bacterial signal peptide sequences. Pfam-scanned protein domain structures are illustrated for each leaf.
